# Obesity-instructed *TREM2^high^* macrophages identified by comparative analysis of diabetic mouse and human kidney at single cell resolution

**DOI:** 10.1101/2021.05.30.446342

**Authors:** Ayshwarya Subramanian, Katherine Vernon, Yiming Zhou, Jamie L. Marshall, Maria Alimova, Fan Zhang, Michal Slyper, Julia Waldman, Monica S. Montesinos, Danielle Dionne, Lan T. Nguyen, Michael S. Cuoco, Dan Dubinsky, Jason Purnell, Keith Heller, Samuel H. Sturner, Elizabeth Grinkevich, Ayan Ghoshal, Astrid Weins, Alexandra-Chloe Villani, Steven L. Chang, Orit Rosenblatt-Rosen, Jillian L. Shaw, Aviv Regev, Anna Greka

## Abstract

Mouse models are a tool for studying the mechanisms underlying complex diseases; however, differences between species pose a significant challenge for translating findings to patients. Here, we used single-cell transcriptomics and orthogonal validation approaches to provide cross-species taxonomies, identifying shared broad cell classes and unique granular cellular states, between mouse and human kidney. We generated cell atlases of the diabetic and obese kidney using two different mouse models, a high-fat diet (HFD) model and a genetic model (BTBR *ob/ob*), at multiple time points along disease progression. Importantly, we identified a previously unrecognized, expanding *Trem2^high^* macrophage population in kidneys of HFD mice that matched human *TREM2*^high^ macrophages in obese patients. Taken together, our cross-species comparison highlights shared immune and metabolic cell-state changes.

## Introduction

Mouse models are an important experimental model organism for studying human diseases, yet differences at the molecular and cellular level between mice and humans complicate the interpretation of mouse-derived data and their translation to the clinic. During kidney development, mice generate around 13,000 nephrons, the functional units of the kidney, over approximately two weeks of active nephrogenesis, whereas human kidneys generate a million nephrons over a 30-week period. In disease, both mouse and human kidneys are known to undergo profound changes in cellular composition (Rennke and Denker, 2019). However, the reasons why some mouse models do not develop kidney disease with the severity observed in humans remain elusive.

Historically, instructive comparisons of corresponding mouse and human tissues have been challenging. On the one hand, high-resolution comparisons of matching cell types require prior knowledge of corresponding markers (which in turn are often fast evolving; (Shay et al., 2013)). Conversely, comparisons based on genomic studies are susceptible to misinterpretation due to averaging over cellular mixtures that either miss fine differences, for example in rare cell types, or obscure similarities in intrinsic cell states due to compositional differences in cell proportions. Single-cell genomics has opened the possibility of a systematic comparison of the cellular composition and gene programs between human and mouse organs. For example, a recent study in human and mouse cerebral cortex (Hodge et al., 2019), showed excellent conservation of many neuronal subtypes, whereas comparisons of the mouse, macaque and human retina (Peng et al., 2019) showed that some neuron subsets are strongly conserved whereas others are difficult to match across species. In this latter case, expression of human disease genes in mouse cell types would lead to erroneous conclusions. Single-cell genomics has been used to chart rich cellular taxonomies of both the mouse (Park et al., 2018) and human kidney (Sivakamasundari et al., 2017; Stewart et al., 2019), including several disease states (Kirita et al., 2020; Wilson et al., 2019; Young et al., 2018). However, studies to date have not tackled a high-resolution side-by-side comparison of mouse and human kidney in health and disease.

With the global obesity epidemic (Pillon et al., 2021), the resulting high rates of type 2 diabetes drive chronic injury and damage to many organs, including the kidney. We focused on diabetic kidney disease (DKD), or diabetic nephropathy, a devastating complication affecting at least 1 in 3 diabetics (https://www.cdc.gov/diabetes). While DKD is the leading cause of kidney failure worldwide (Jager et al., 2019), many questions remain unanswered. First, little is known about kidney cell states associated with high fat diet exposure at the earliest stages of disease, before overt organ damage. Second, DKD pathophysiology is inherently complex, involving hemodynamic, metabolic and immune factors. Growing evidence suggests that chronic, low-grade inflammation plays a key role in the progression of DKD (Wada and Makino, 2016) but the role of individual immune cell types in exacerbating or ameliorating disease is an ongoing area of investigation. Third, the relative paucity of kidney biopsies from patients with DKD has hindered deep mechanistic insights and has further heightened the need for reliable animal models. However, to date, there has been no agreement on a mouse model that incorporates salient features of human DKD (Betz and Conway, 2014). Assessing the potential of mouse models to recapitulate human DKD at the cellular level, with a focus on the earliest immune and metabolic changes, would expand our fundamental understanding of disease mechanisms, and demonstrate advantages and limitations of specific mouse models for therapeutic discovery. Most importantly, despite recent advances that help manage and mitigate disease progression (e.g. SGLT2 inhibitors; (Ingelfinger and Rosen, 2019; Perkovic et al., 2019)), curative therapies for DKD are still urgently needed. Therefore, understanding the earliest steps connecting obesity, diabetes and kidney injury may reveal a time-window and opportunity for curative therapeutic interventions.

Here we compared and contrasted the regional and cellular states of the healthy adult kidney in mouse and human, identifying shared broad cell classes alongside species-specific granular states. We observed macrophage heterogeneity in both species, and describe *LYVE1^high^* and *TREM2^high^* macrophages in the human adult kidney resembling subsets in other tissues. Next, we collected matching single-cell RNA-seq profiles of the kidney from two mouse models of DKD and from several patients. In particular, we probed for progressive changes in kidney cell populations in a high-fat diet (HFD) model and a genetic model (BTBR *ob*/*ob*) of DKD, at multiple time points along disease progression. By comparing this rich mouse dataset with kidney tissue from obese and diabetic patients in the early stages of disease, we identified obesity-instructed *Trem2^high^* macrophages as a shared cellular feature of kidney injury in mouse and human.

## RESULTS

### Matching single-cell RNA-Seq of mouse and human kidney

To systematically compare cellular taxonomies of the mouse and human kidney, we generated regional scRNA-seq maps in each of three anatomical regions: cortex, medulla, and hilum (pelvis), with a consistent regional sampling strategy in both species (**Figure 1A, B**). For human, we dissociated macroscopically normal tissue from tumor nephrectomies of 9 donors and recovered 76,868 scRNA-seq profiles (**Methods, Table S1, Figure 1**), spanning 30,072 cortical, 26,949 medullary, and 19,847 hilar cells. For mouse, we macroscopically dissected 10-12-week-old male and female mouse kidneys (**Figure 1B**), and obtained 148,647 scRNA-seq profiles, spanning 43,246 cortical, 51,437 medullary and 53,964 hilar cells, with another 73,214 cells derived from coronal sections (**Methods, Table S1,Figure S1**).

**Figure 1.**
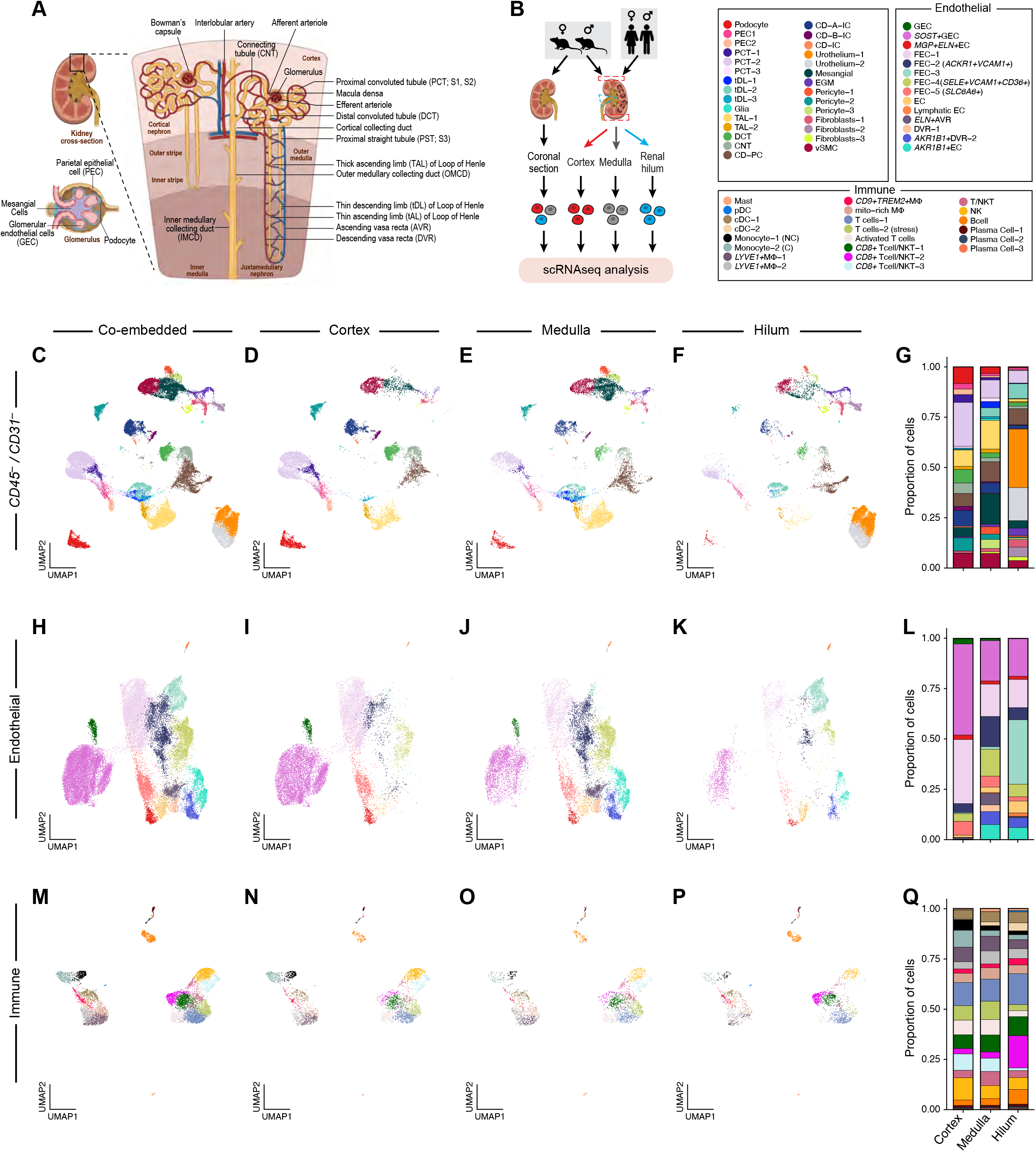
Regional sectioning of human kidney reveals spatial enrichment for distinct kidney cell subsets by scRNA-seq. (A) Graphical representation of kidney anatomy. Each functional unit or nephron consists of the glomerulus within Bowman’s capsule and tubule. (B) Schematic depicting the scRNA-seq study design for the mouse and human regional transcriptomic atlases. Sampled regions include: cortex (red), medulla (gray) and renal hilum (blue). For mice, coronal sections were also included. (C-G) Comparison of major human kidney *CD45*-/*CD31*-cell subsets across three regions. Uniform manifold approximation and projection (UMAP) visualization of cell subsets obtained via co-embedding (C) of the different kidney regions: cortex (D), medulla (E) and renal hilum (F). Points represent individual cells and colors represent cell subsets shown in the legend. (G) Barplot shows the proportion of cells belonging to each cell class by sampled region. PEC, parietal epithelial cells; EGM, extraglomerular mesangial cells; PCT, proximal convoluted tubule; tDL, thin descending limb; TAL, thick ascending limb; DCT, distal convoluted tubule; CNT, connecting tubule; collecting duct cells: CD-PC, principal; CD-A-IC, intercalated alpha; CD-B-IC, intercalated beta; vSMC, vascular smooth muscle cells. (H-L) Comparison of human kidney endothelial (*CD31*+) cell subsets across three regions. UMAP visualization of cell classes obtained via co-embedding (H) of the different kidney regions: cortex (I), medulla (J) and renal hilum (K). (L) Barplot shows the proportion of cells belonging to each cell class by sampled region. GEC, glomerular endothelial cells; FEC, *PLVAP*+ fenestrated endothelial cells; DVR, descending vasa recta; AVR, ascending vasa recta; LEC, Lymphatic endothelial cells. (M-Q) Major immune cell classes were recovered from all human kidney regions. UMAP visualization of cell classes obtained via co-embedding (M) of the different kidney regions: cortex (N), medulla (O) and renal hilum (P). (Q) Barplot shows the proportion of cells belonging to each cell class by sampled region. pDC, plasmacytoid dendritic cells; cDC, conventional dendritic cells; Monocyte-1 (NC), Non-Classical; Monocyte-2 (C), Classical; NK, Natural Killer; NKT, Natural Killer T cell.

We partitioned cells by the expression of known markers into three groups: endothelial (*Pecam1*^+^/*PECAM1*^+^/*CD31*^+^) (**Figure 1H-K, Figure S1F-J**), immune (*Ptprc*^+^/*PTPRC*^+^) (**Figure 1M-P, Figure S1K-O**), and the remaining *CD31_-_*/*CD45*^-^ group comprised of epithelial, stromal and other cells types (**Figure 1C-F, Figure S1A-E, Methods, Table S2**). We then iteratively clustered the cells in each of these groups (separately for human and mouse to accommodate species-specific genes, and combining all regions), first into major kidney cell types (“broad cell classes,’’ e.g. proximal tubular cells (PCT)) and then into finer, higher resolution subsets (“granular subsets,” e.g. PCT-1, 2, 3, etc.), and annotated each *post-hoc* by differentially expressed genes and known canonical markers (Stewart et al., 2019; Subramanian et al., 2019; Young et al., 2018) (**Figure S2, S3A-B, Table S2-4**).

We found concordance in cellular composition between mouse and human for all three kidney regions (**Figure 1G, L, Q; Figure S1P, Q, R**) as expected by anatomy (Rennke and Denker, 2019). For example, podocytes (human cortex: 8.3%, medulla: 3.2%, hilum: 0.5% vs. mouse cortex: 4.7 %, medulla 1.6 %, hilum 3.8%), glomerular parietal epithelial cells (PECs) (human cortex: 5.4%, medulla: 2%, hilum: 1.2% vs. mouse cortex 2.1%, medulla 0.47%, hilum 1.1%), and proximal tubular cells (PCT) (human cortex: 26.6%, medulla: 11.9%, hilum: 6.4% vs. mouse cortex: 29.5%, medulla 13%, hilum 18.9%) were enriched (p value 3.93e-04 (human), <2e-16 (mouse)) in kidney cortex. To facilitate future studies, we provide a searchable online resource (https://singlecell.broadinstitute.org/single_cell) for the expanded kidney cellular taxonomies.

### Over 60 human kidney cell subsets identified, with regional enrichment and including rare cell types

In the adult human kidney, we identified 32 broad cell classes (e.g. PCT, podocytes) across 65 granular subsets (e.g. PCT-1, 2, 3, etc.) (**Figure S2A,C, Figure S3A, Table S2)**, encompassing the major expected parenchymal cells of the human kidney nephron, and multiple subsets of fibroblasts, pericytes including a distinct *AGTR1*+ subset, and urothelial cells (the latter found almost exclusively in the hilum, the region closest to the ureter/bladder). We also annotated major immune cell populations including macrophages, B, T, Natural Killer (NK), mast and dendritic cells (DC) across all well-described regions (Stewart et al., 2019) noting multiple subsets of macrophages, and *CD8*+ and *CD4*+ T or NKT (Natural Killer T) subsets (**Figure S3A, Table S3**). Finally, turning our attention to rare cell types, we identified cells of the macula densa expressing *NOS1*, as well as glial cells expressing *S100B*^+^, *PLP1*^+^ and *CDH19*^+^ (**Table S3**).

### Newly identified LYVE1^high^ and CD9^high^TREM2^high^ macrophages in adult human kidney

By iterative clustering, we identified five macrophage subsets that have not been previously described in the human adult kidney (**Methods, Figure 2A, Table S4**): three *LYVE1^high^* subsets (*LYVE1^high^*-1(*CCL3^low^CXCL1^low^*) *LYVE1^high^*-2 (*CCL3^high^CXCL1^low^*), *LYVE1^high^*-3 (*CCL3^low^CXCL1*^high^), and two *CD9^high^* subsets (*CD9^high^TREM2^low^*, *CD9^high^TREM2*^high^). The *LYVE1^high^* subsets were characterized by high expression of genes associated with homeostatic macrophage populations (Cochain et al., 2018; Pinto et al., 2012), including *SEPP1*, *FOLR2*, *LGMN*, *CST3* and *DAB2* (**Figure 2A**), in line with the LYVE1^high^CX3CR1^low^ tissue interstitial macrophages recently described in human lung, adipose, and multiple mouse tissues (heart, fat, and skin; Chakarov et al. 2019). *LYVE1^high^* kidney macrophages expressed all markers also found in *LYVE1^high^* heart macrophages (Litviňuková et al. Nature 2020) (**Figure S3C**). The *LYVE1^high^*-2 and *LYVE1^high^*-3 subsets were distinguished from the *LYVE1^high^*-1 subset by expression of specific chemokines (*CCL3*, *CCL4*, *CCL4L2*, *CCL2*), suggestive of an activated state, similar to previously described senescent-like microglia expressing inflammatory genes (Geirsdottir et al., 2020).

**Figure 2.**
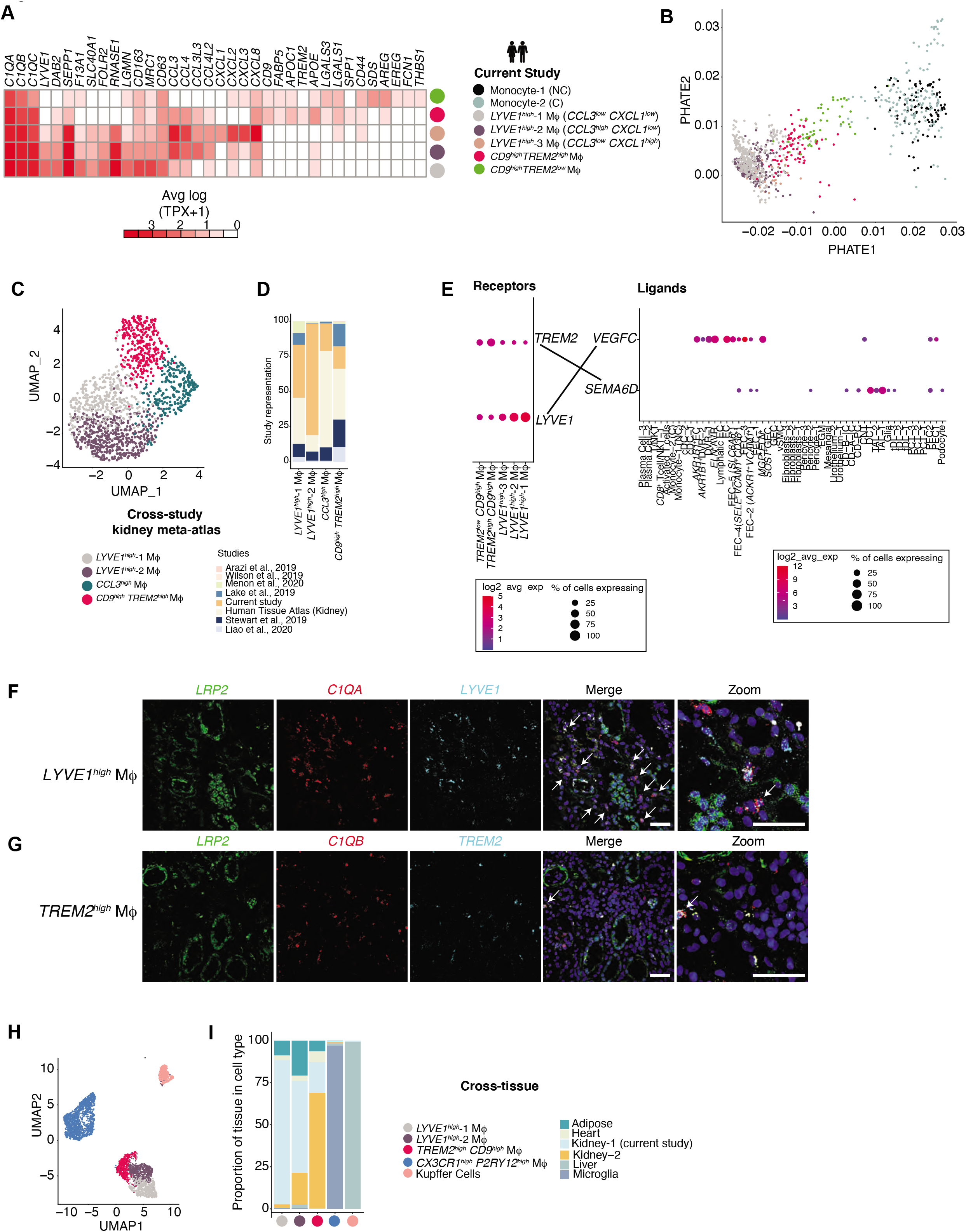
Heterogeneous macrophage subsets in human kidney at single cell resolution. (A) Heatmap visualization of expression programs distinguishing the 5 macrophage (MΦ) populations in the human adult kidney: *LYVE1*+, *CCL3*+*LYVE1*+, *CXCL1*+*LYVE1*+, *CD9*+*TREM2*- and *CD9*+*TREM2*+. Columns are genes that are differentially expressed in populations. *C1QA* and *C1QB* are pan MΦ markers. Values (color) represent column-normalized average gene expression in units of log(TPX+1). (B) Potential of Heat diffusion for Affinity-based Transition Embedding (PHATE) visualization of macrophages (MΦ) in the human kidney along with classical and non-classical monocytes. Data shown from 3 nephrectomies. (C) UMAP visualization of MΦs from kidney meta-atlas derived from 7 different studies (D) Barplot of study representation among the cross-study macrophage subsets. (E) Selected receptor-ligand interactions. (F,G) Characterization of *LYVE1*+ (F) and *TREM2+CD9*+ (G) macrophage subsets by *in situ* HCR; arrows identify individual cells. (H) UMAP visualization of co-embedding of MΦs from human adipose, heart, liver, microglia and kidney reveals 5 populations: *LYVE1*+, *CCL3*+*LYVE1*+, *CX3CR1*+*PY2R1*+, *CD9*+*TREM2*+, and Kupffer cells. (I) Barplot of tissue composition among the cross-tissue macrophage subsets.

While both *CD9^high^* populations shared transcriptional signatures (**Figure 2A**, e.g., *CD9*, *FABP5*, *APOE*, *APOC1*, *LGALS3*, *LGALS1*), *CD9^high^TREM2^high^* cells additionally showed induction of *TREM2, SPP1*, and cathepsins, reminiscent of the Lipid (or Scar)-Associated-Macrophage (LAM/SAM) signature described in human adipose tissue and liver in the setting of inflammation (Hill et al., 2018; Jaitin et al., 2019). Since *CD9^high^TREM2^high^* cells are suggested to be monocyte-derived (Jaitin et al., 2019; Ramachandran et al., 2019), we examined kidney macrophage profiles by Potential of Heat diffusion for Affinity-based Transition Embedding (PHATE, (Moon et al., 2019)). The newly identified macrophage subsets lay along a spectrum spanning monocytes, *CD9^high^TREM2^low^* macrophages, *CD9^high^TREM2^high^* (LAM-like) macrophages, and different subsets of homeostatic *LYVE1^high^* macrophages (**Figure 2B**). This analysis suggested that these newly described kidney *CD9^high^TREM2^high^* cells were transcriptionally more similar to monocytes than to homeostatic macrophages.

All five macrophage subsets were present in the 9 human donors (**Table S4**), and also identified in a recently assembled kidney meta-atlas (Subramanian et al., 2020), **Figure 2C-D, Figure S3D**). In the context of cell-cell interactions, *VEGFC*, a putative LYVE1 ligand, was expressed in PECs and connecting tubule cells (CNT) at greater than 5% prevalence in addition to its expected expression in endothelial cells (ECs) (**Figure 2E)**. Among several reported TREM2 ligands (Kober and Brett, 2017), only *SEMA6D*, expressed in podocytes, PECs, glia, and a number of distal tubular epithelial cells (tDL, DCT, TAL, CD-PC, CD-A-IC) crossed the prevalence threshold. We confirmed that *LYVE1^high^* and *TREM2^high^* macrophages localized to the tubulointerstitial compartment of adult human kidney using *in situ* hybridization (**Figure 2F-G**), demonstrating spatial context (with PCT marker *LRP2*) and using the pan-macrophage markers *C1QA* or *C1QB*, the homeostatic macrophage marker *FOLR2* for *LYVE1^high^* and *LGALS* for *TREM2^high^* populations, respectively (**Figure S4A-B**).

We next asked if the macrophage states we identified in the human kidney are present in other human tissues. Co-embedding our kidney macrophage profiles with macrophages from scRNA-seq studies of adipose tissue, kidney, heart, liver, and brain (microglia) (Han et al., 2020; Keren-Shaul et al., 2017), resulted in five macrophage populations (**Figure 2H-I**; **Methods**). These included 3 *TREM2*^high^ populations: *CX3CR1*^high^*P2RY12*^high^ microglia; *CD5L^high^MARCO^high^TREM2^high^* liver Kupffer cells and the *CD9^high^TREM2^high^* macrophages shared across adipose tissue, heart and kidney. The *LYVE1^high^* and *CCL3^high^LYVE1^high^* populations were present in kidney, liver, as well as heart and adipose tissue (Chakarov et al., 2019). We concluded that the distinct *LYVE1^high^* and *TREM2^high^* macrophage populations we identified in human adult kidney shared signatures across several tissues.

### The coronal section captures cell states found in all three regions of the adult mouse kidney

Turning our attention to the mouse kidney, we identified 35 broad cell classes and 87 granular subsets (**Figure S2B, D, Figure S3B, Table S5**). While the cortex, medulla, and hilum included region-specific subsets, we found that the coronal section effectively captured all 87 subsets, suggesting that coronal sections are sufficiently representative of the entire mouse kidney. Of interest, we recovered three subsets of mouse PECs from the cortex, including a previously uncharacterized PEC-1 population expressing neuronal genes (*Pcp4*, *Tnc*), podocyte developmental markers (*Wt1*, *Ncam1*), and smooth muscle markers (*Myl9*, *DKk3*) (**Figure S4C**), and validated them as PECs by *in situ* hybridization (**Figure S4E, F**), based on their localization along the periphery of podocyte-lined glomeruli (Kuppe et al., 2015, Kuppe et al., 2019). Additionally, we validated PCT segment-specific markers predicted by our analysis, by *in situ* hybridization, including SGLT1 (*Slc5a1*), SGLT2 (*Slc5a2*), *Slc22a8*, *Slco1a1 and Acsm3* (**Figure S4D, S4G-I**). Our data confirmed previously described sexual dimorphism in the mouse PCT (Ransick et al., 2019) (**Figure S5A-B**).

Our regional enrichment strategy for the mouse revealed several rare cell types. For example, iterative clustering of cortical mesenchymal and medullary thick ascending limb (TAL) cells, respectively, identified cellular components of the juxtaglomerular apparatus (JGA), a rare heterogeneous population of cells, including renin-secreting cells (*Ren1*^+^), and 26 cells of the macula densa, involved in the regulation of blood pressure and glomerular filtration rate (Peti-Peterdi and Harris, 2010) (**Methods**, **Figure S5C-J**). We confirmed the JGA and macula densa cells by *in situ* hybridization for *Ren1* and *Nos1* respectively and the macula densa signature gene *Ptgs2* (**Figure S5E, Figure S5J**). We also identified rare glial cells in mouse kidney, significantly enriched (cortex: 0.02%, medulla 0.04%, hilum 1.5%, p-value < 2e-16) in the hilum, consistent with a role in innervation (Sata et al., 2018) (Figure S1P). Finally, we identified rare type 2 innate lymphoid cells (ILC2) (**Figure 1M-Q**; **Figure S1K-O,R**) in the hilum.

The macrophages in mouse were clearly partitioned by recently reported signatures of mouse ‘resident’ (*C1q*, *ApoE*, *Cd74*, *Ms4a7*, *Lgmn*, *Hexb*, *Cxcl16*) and ‘infiltrating’ (*Ccr2*, *Ifitm6*, *Chil3*, *Ear2*, *Crip1*, *S100a4*, *Plac8*) macrophages (**Figure S3B**)(Zimmerman et al., 2019). *In situ* hybridization using markers for resident macrophages (*C1qa*, *C1qb*) and glomerular (*Nphs2*; podocin) or PCT markers (*Lrp2*; megalin), localized macrophages adjacent to glomeruli, along with *Siglech*^+^ pDCs and *Naaa*^+^ cDCs (**Figure S6A-I**) near fenestrated endothelial and lymphatic vessels. Among mouse macrophages, distinct populations with differential gene expression of *Retnla* (resistin-like alpha), *Mrc1* (mannose receptor, CD206), and *Mgl2* (macrophage galactose N-acetyl-galactosamine specific lectin 2) were identified as homeostatic resident macrophages (Pinto et al., 2012). Enriched at the renal hilum (**Figure S1N**), they also differentially expressed *Lyve1* and other “M2” genes such as *Folr2, Sepp1* and *Igf1* (Pinto et al., 2012). The hilar *Lyve1^+^* population also expressed *Mmp9*, recently found to be instrumental in the prevention of collagen deposition by vascular smooth muscle cell (vSMC)-associated *Lyve1*^+^ macrophages (Lim et al., 2018). Thus, our analysis identified multiple subsets of macrophages in the mouse kidney with distinct regional distribution patterns.

### Broad kidney cell classes are conserved between mouse and human with species-specific granular cell subsets

We compared the diversity of cell classes recovered from mouse and human kidney. Except for mast cells, which we only recovered from the human kidney, and the thin ascending limb (tAL), ILCs and neutrophils, which we could only definitively discern in the mouse kidney, the majority of broad cell classes (30, human=90.1%, mouse=85.7%) were shared between the two species with some differences in regional distribution (**Table S6**): mouse lymphatic endothelial cells (LECs) and glia were only recovered from the hilum, whereas human LECs, distinguished by *LYVE1*, *MMRN1*, and *ACKR2*, and glia spanned all kidney regions. While both mice and humans had multiple fibroblast and pericyte populations, we observed a discrete population of human *AGTR1*^+^ pericytes. Finally, we recovered more granular subsets (87 vs 65) from mice, possibly due to deeper sampling.

To determine the extent of similarity in cell type specific signatures, we used a multi-class random forest classifier trained on mouse cell subsets to classify human cells (Peng et al., 2019), and vice versa (**Methods**, **Figure S7A-C**) at two levels of annotation: broad cell classes and granular subsets. The specificity of the mapping was quantified as (a) 1-1 if greater than 90% of the cells in the human subset mapped to the mouse subset, (b) orthologous and highly specific if 80-90% mapped (c) orthologous if 65-80% mapped and (d) non-specific if <65% mapped (**Table S6**).

Broad cell classes (**Figure 3A,D,G**) had high inter-species 1: 1 correspondence, but subsets (**Figure 3B,E,H**) were more divergent. Among broad classes, epithelial cells had high mouse-human correspondence ranging from 98% mapping for the thick ascending limb (TAL) to 92.2% for podocytes (suggesting strong conservation of marker genes). In contrast, mesenchymal cells had on average lower correspondence of 11.4%, potentially due to shared signatures between mesenchymal classes, and hence, cross-mapping. At the granular level, mapping of PCT and thin descending limb (tDL) subsets was diffuse, whereas mesenchymal cells showed more specificity. Among rare cells, we found that, while both mouse and human macula densa cells were *Nos1*^+^/*NOS1*^+^, their gene signatures were distinct (**Figure S5G, S5I**).

**Figure 3.**
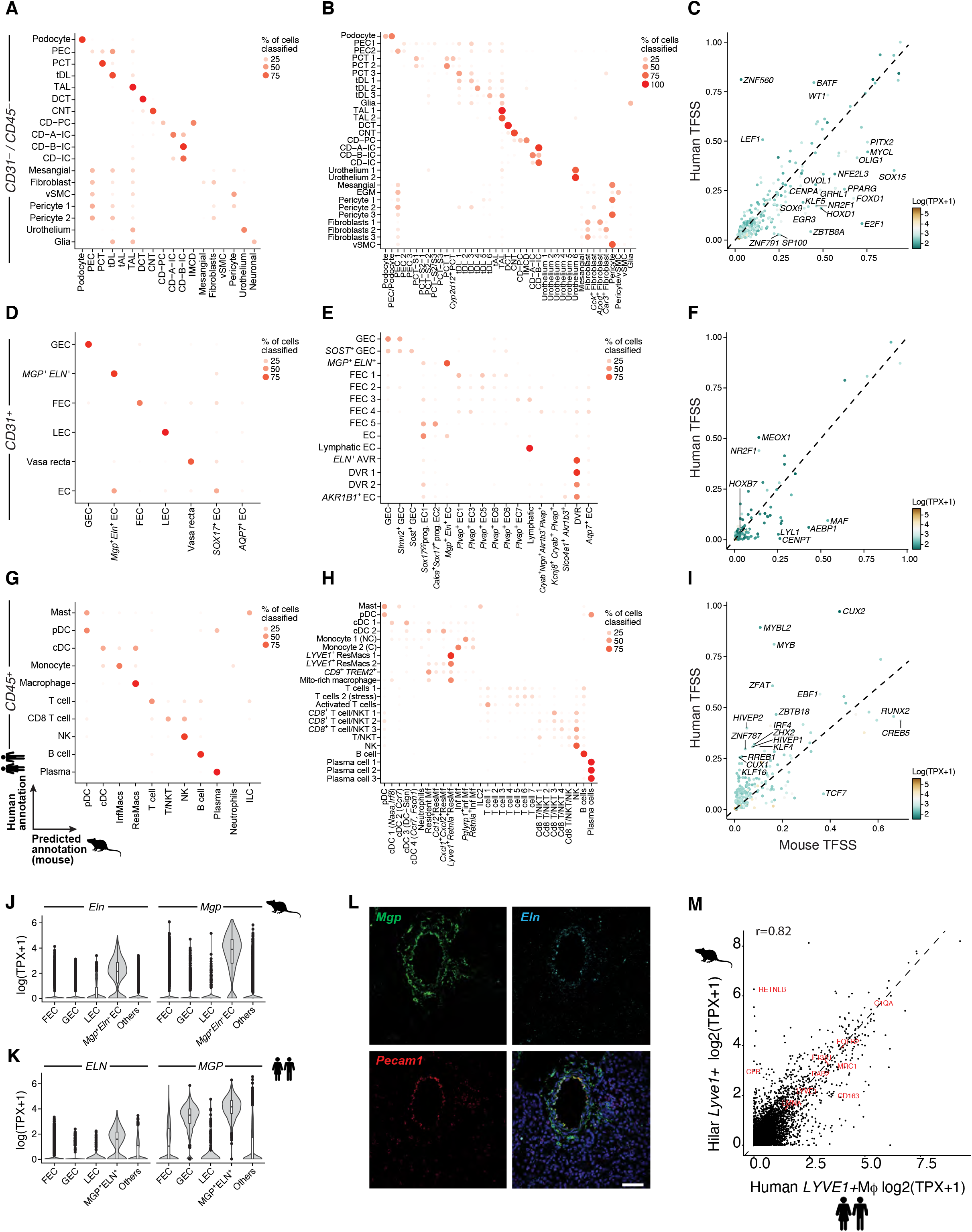
Shared and distinct cell-type specific transcriptional programs in mouse and human kidney. (A,B) Dot plots representing prediction results in human data of random-forest classifier trained on mouse Cd45-/Cd31- (A and B), cells. The size and color of the dot represents the proportion of cells in a given human cell type (Y-axis) that was assigned a cell type label corresponding to or classified as the mouse cell type (X-axis). (C) Comparison of transcription factor cell type specificity between mouse and human in the Cd45-/Cd31-compartments. (D, E) Dotplot representation of cell-type specific human transcriptional program (rows) scores in mouse kidney Cd31+ cells (columns). A random sample of 5 cells per mouse cell type are shown for brevity. Scores are row-normalized. (F) Comparison of transcription factor cell type specificity between mouse and human in the Cd31+ compartments. (G, H) Dotplot representation of cell-type specific human transcriptional program (rows) scores in mouse kidney Cd45+ cells (columns). A random sample of 5 cells per mouse cell type are shown for brevity. Scores are row-normalized. (I) Comparison of transcription factor cell type specificity between mouse and human in the Cd45+ (G) compartments. (J) Violin plots reveal a novel Mgp+Eln+ subset in mouse and human endothelial cells. (K) Immunostaining of Mgp and Eln with Pecam1.

In the endothelial cell compartment, GECs (**Figure S7D-E**) and LECs (**Figure S7F-G**) mapped 1:1, while *MGP*^+^*ELN*^+^EC had 87% correspondence. *MGP^+^ELN*^+^EC (**Figure 3J-K**), coexpressing *Mgp*, a matrix Gla protein known for its involvement in endothelial-epithelial interactions (Yao et al., 2016), and elastin (*Eln*), known to regulate vessel stiffness in mice (Wagenseil and Mecham, 2012) was found in all regions in both species. In situ hybridization (**Figure 3L**) in mice revealed *Mgp*^+^ *Eln*^+^EC were associated with vessels larger than GEC (**Figure S7D-E**) or FEC (**Figure S7D-E**), suggestive of interlobular vessels. All regions in mice shared a distinct *Sox17*-expressing (*Sox17*^+^) progenitor population with a subset expressing the angiogenesis promoter calcitonin (*Calca*), grouped into the broad “*Sox17*^+^” class, whereas in human, *SOX17* was expressed broadly across endothelial classes.

Among immune cells, NK, B, plasma cells and macrophages had strongly conserved markers. Some granular macrophage subsets, such as *LYVE1^high^*MΦ-1 mapped 1:1 to their mouse counterparts, while others, such as monocyte-1 (classical) cells and *CD9^high^TREM2^high^* MΦs did not specifically map to any one of the mouse infiltrating or resident subsets. The diffuse mapping of granular subsets may reflect species-specific cell states (*e.g*., TAL-2), or limitations of discrete clustering, applied independently to each species (*e.g*., Fibroblast-1, PCT-1). Even within classes with 1-1 correspondence such as *LYVE1^high^* macrophages, species-specific marker genes may be present (**Figure 3M**). Our comparisons emphasize that while broad identity programs are shared between mouse and human, it is important to understand which granular subsets and cell states may be shared.

### Conservation of TF cell specificity between human and mouse

We hypothesized that conserved cell classes may be established, in part, by conserved Transcription Factor (TF) regulatory mechanisms, which may be reflected by the cell-type specificity of TF expression between the two species. To this end, we computed TF specificity scores (TFSS) (Peng et al., 2019) in each of human and mouse restricting to TFs with a certain minimum prevalence, based on the Shannon diversity index (**Methods**) for the entire shared set of broad cell classes (a total of 30, **Table S6**). Low TFSSs (high Shannon indices) should correspond to an even distribution of the expression of TFs across subsets, as one would expect for constitutively expressing TFs (e.g. *HOXB7* in endothelial cells, **Figure 3F**). Conservation of TF specificity should be reflected in correlation of TFSS profiles. Human and mouse TFSS were correlated in all three compartments (**Figure 3C, F, I;** r=0.82, 0.59, 0.59, p<1e-16). Within the *CD31*^-^/*CD45*^-^ cells, the expression pattern of certain TFs relevant to kidney homeostasis such as *SREBF2* (lipid metabolism) (Dorotea et al., 2020), *MLX* (glucose metabolism) (Suzuki et al., 2020), *TCF7L2* (associated with CKD function) (Köttgen et al., 2008) were highly congruent between species (**Figure 3C**). While *FOXD2* was conserved with specificity to podocytes in both species (TFSS hs=0.51, mm=0.5), *FOXD1* was less specific in humans when compared to mouse podocytes (TFSS hs=0.2, mm=0.67). On the other hand, *KLF2* (TFSS hs=0.08, mm=0.082) had low cell type specificity in both species. Overall, we noted higher inter-species TFSS concordance in the epithelial cell compartment (r=0.82), compared to the other compartments.

In the endothelial compartment (**Figure 3F**), most TFs had low cell type specificity in both mouse and human. NoTable exceptions were *PROX1* (TFSS hs=0.81, mm=0.77), *TBX1* (TFSS hs=0.98, mm=0.91) and *FOXP2* (TFSS hs=0.87, mm=0.96) that were specific to LECs, while *LEF1* (TFSS hs=0.79, mm=0.64) was specific to the vasa recta (VR) in both species. Among divergent TFs, *Maf* (TFSS hs=0.79, mm=0.64) and *Aebp1* (TFSS hs=0.79, mm=0.64) were specific to mouse LECs, while *MEOX1* (TFSS hs=0.79, mm=0.64) and *NR2F1* (TFSS hs=0.44, mm=0.14) was specific to human fenestrated endothelial cells (FECs). Similarly, within the immune compartment (**Figure 3I**), *MYB*, *MYBL2* and *CUX2* (human pDCs) had higher TFSS in humans than in mice, whereas *TCF-7* TFSS was higher in mouse (specific to T-cells). *LEF1* was conserved and specific to T-cells in both species. TFs whose TFSS’s deviated between species may be indicative of differences in activity between the two species. For example, *SOX15*, which has a much higher TFSS in mouse, was broadly expressed in human parenchymal cells, whereas expression was restricted to specific subsets in mice. Thus, differences in transcriptional regulation within cell states between species may play a role in species-specific cell states.

### Comparative analysis of disease states associated with obesity and diabetes in mouse and human kidneys

Turning our attention to obesity- and diabetes-driven kidney disease (DKD), the most prevalent cause of kidney failure in humans (Maric-Bilkan, 2013), we sought to profile kidneys from two mouse models. Specifically, we analyzed a metabolic model of high-fat diet (HFD)-induced kidney injury and a well characterized genetic model of *ob*/*ob* leptin deficiency on a BTBR background (Hudkins et al., 2010). We compared the single cell profiles derived from these mice to kidney nephrectomy tissue from patients with obesity and histologically evident, early signs of DKD (**Figure 4A**). We focused on cortical samples, because, based on regional scRNA-seq studies (**Figure 1**), we concluded that cortical sections can sufficiently capture most kidney cells, while significantly reducing time and cost.

**Figure 4.**
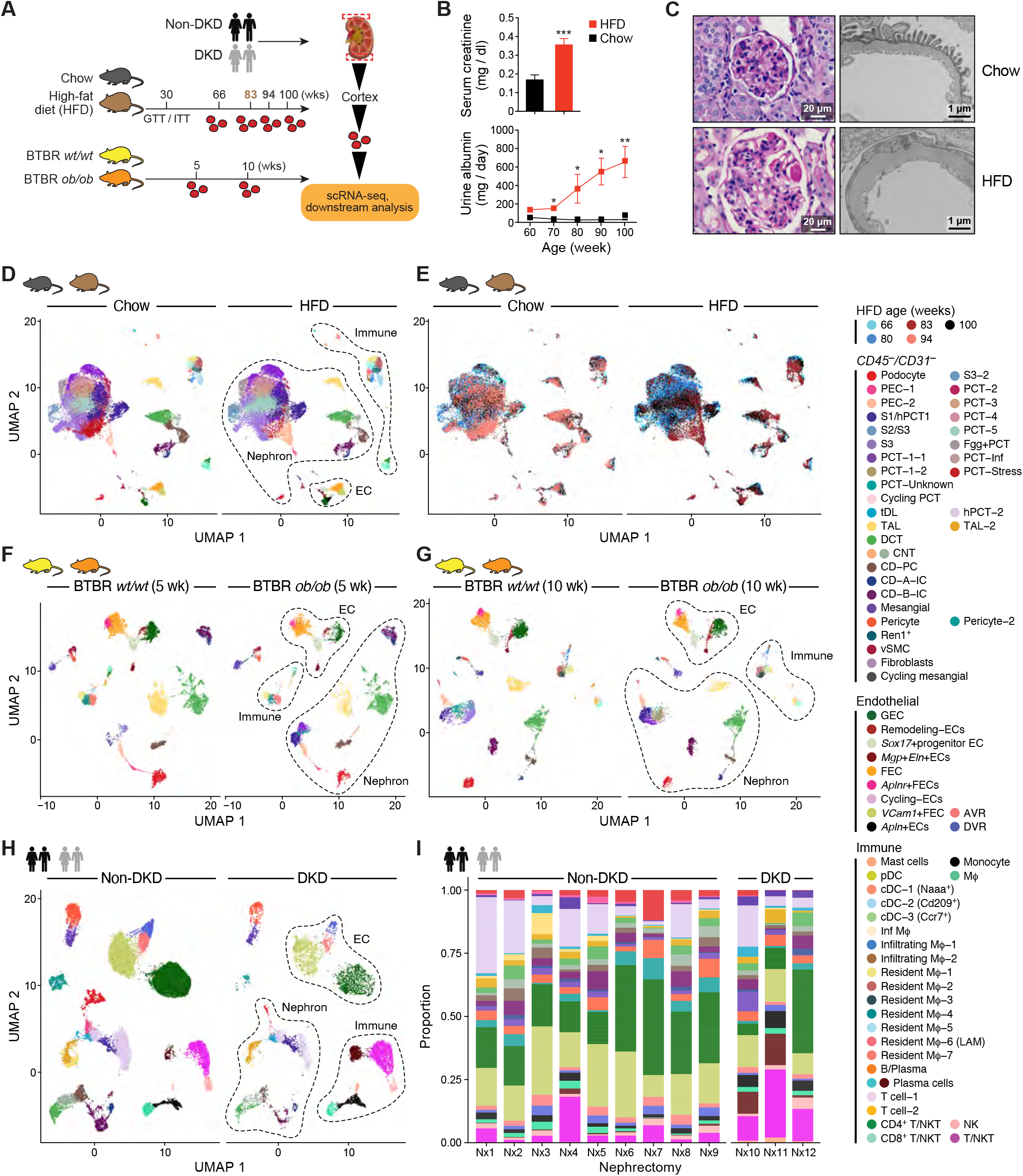
Single-cell atlases of diabetic kidney disease in mouse and human. (A) Experimental timeline and sampling outline for kidney tissue from two mouse models and nephrectomy specimens from patients with and without DKD. Glucose and insulin tolerance tests were performed at 30 weeks of age in the HFD mouse model, and in the week prior to tissue collection in the 5- and 10-week-old BTBR mice. Kidney cortex obtained at equivalent time points from HFD- and chow-fed mice (aged 66-100 weeks), BTBR *ob*/*ob* and BTBR *wt*/*wt* mice (aged 5-10 weeks) and patients undergoing tumor nephrectomy, was dissociated using the same enzymatic protocol. Single cells from each sample were loaded into individual channels of a 10x Genomics Chromium 3’ chip. (B) Phenotypic characterization of the HFD-fed mice including serum creatinine concentration and albuminuria quantification, *p<0.05, **p<0.01, ***p<0.001. Serum creatinine chow (n = 32), HFD (n = 37), p<0.0001; urine albumin for chow (n = 57) and HFD (n = 106). (C) Light and transmission electron microscopic examination of kidney tissue from chow and HFD-fed mice. (D) UMAP plots of cells recovered from the kidneys of chow- and HFD-fed mice. Individual populations are represented by distinct colors. (E) UMAP plots of the cells from kidneys of chow- and HFD-fed mice, colored according to mouse age. (F) UMAP plots of cell populations identified in the kidneys of 5-week-old BTBR *wt*/*wt* and BTBR *ob*/*ob* mice. Individual populations are represented by distinct colors. (G) UMAP plots of cells recovered from the kidneys of 10-week-old BTBR *wt*/*wt* and BTBR *ob*/*ob* mice. Individual populations are represented by distinct colors. (H) UMAP plots of the cell populations identified in nephrectomy specimens from patients with and without DKD. Individual populations are represented by distinct colors. (I) Relative cell proportions in each patient nephrectomy sample. Individual populations are represented by distinct colors.

First, we characterized both mouse models to confirm the onset and progression of kidney injury (**Figure 4B-C, Figure S8M, Q)**. 10-week-old C57BL/6-129 mice were fed a HFD for more than 90 weeks, alongside littermate controls fed a normal chow diet (**Figure 4A-B**). We confirmed weight gain, insulin resistance and kidney dysfunction at 20 and 60 weeks, respectively (**Figure 4B, S8A, C**). This was driven by significant obesity, dyslipidemia, and hormonal dysregulation, as has been previously described (Yore et al., 2014) (**Figure S8A, D-I**). Most importantly, HFD-fed mice developed progressive kidney disease, with marked kidney filter damage (demonstrated by increased urinary albumin or albuminuria; **Figure 4B, lower**) and elevation in serum creatinine at the terminal time point (~90 weeks; **Figure 4B, upper**). Histologically, HFD-fed mice developed significant glomerular hypertrophy and nodular sclerotic lesions visible by light microscopy (**Figure 4C, left lower**), as well as significant glomerular basement membrane thickening, visualized by electron microscopy (**Figure 4C, right lower**), all hallmarks of DKD in humans.

As a comparator, we studied BTBR *ob*/*ob* mice, known to exhibit kidney failure by 20 weeks of age (Hudkins et al., 2010), which prompted us to study them at two earlier time points, when changes in the kidney reflect actively progressing injury (5 and 10 weeks of age). Consistent with prior reports, and similar to HFD-fed mice, BTBR *ob*/*ob* mice at 5 and 10 weeks developed insulin resistance, marked urinary albumin and glomerular basement membrane thickening as signs of kidney filter damage, as well as significantly elevated cholesterol, but no creatinine elevation (**Figure S7K-Q**).

Finally, out of 12 human nephrectomies, 6 patients were clinically obese, and a subset of three patients had specific histologic evidence of early kidney injury, with diffuse and early nodular diabetic glomerulosclerosis (DKD) on light microscopy determined by an experienced renal pathologist blinded to our study design (**Table S1**). We confirmed podocyte loss in tissue from these DKD patients by in situ hybridization (**Figure S8R, S**).

Our analysis of kidney injury in mouse and human spanned 74,107 cells from the HFD mouse model, 22,696 and 18,686 cells from the BTBR model at 5 and 10 weeks respectively, and 40,468 cells from human samples. Fewer cells were recovered from the BTBR *ob*/*ob* mice compared to the BTBR *wt*/*wt* controls at both time points (**Table S7**). We observed a discrepancy between the proportion of PCT cells captured in the HFD and BTBR mouse models, which may be due to differences between mouse strains and/or sensitivity to the single cell dissociation protocol, favoring recovery of PCT cell types in C57BL/6-129 mice. Overall, broad cell classes were consistent between the two mouse models with 24 shared classes (**Table S7**). Each strain had its own granular subsets, most evident among PCT, endothelial and immune cells. Joint analysis of human non-DKD and DKD samples (we conservatively selected the three samples that had overt histologic evidence of DKD for this analysis) captured parenchymal, stromal and immune cell classes (**Figure 4D-I**), with 15 shared cell classes between both mouse models and humans.

We next asked whether the expression changes associated with DKD in matching cell types were conserved between human and mouse. We identified genes that were differentially expressed in each cell type between DKD and non-DKD samples, while accounting for donor-specific random effects in a regression framework for the human data (**Methods, Table S8**). We prioritized significant genes based on effect sizes after adjustment for false discovery rate (FDR<0.1).

Overall, we found fewer differentially expressed genes in human than in mouse (**Figure S9A-D, Table S8**). In human early DKD, distal tubular cells (DCT, TAL **Figure S9D**) displayed the highest numbers of differentially expressed genes as previously reported (Wilson et al., 2019). The DCT (**Figure S9E**) and TAL (**Figure S9F**) thus showed the most cross-species overlap in disease gene signatures, with shared DE genes enriching for the immunoproteosome, lysosome and phagosome (DCT) and oxidative stress (TAL) pathways. In podocytes and mesangial cells, the transcriptional response to disease varied between mouse and human, both in terms of the specific genes affected and the direction of the effect (**Figure S9G-H**). Such cross-species differences could be due to biological differences in the natural history of disease, high donor-donor variability or experimental limitations (smaller numbers of cells recovered, heterogeneity of sample size, etc.).

Across mouse models at all time points, podocytes and mesangial cells exhibited significant changes in expression profiles, in line with well described roles for these cell types in the progression of diabetic kidney injury (Brunskill and Potter, 2012) (**Figure S9A-C, G-H**). We validated the collagen genes *Col4a3* and *Col4a2*, upregulated in diabetic podocytes and mesangial cells, respectively, by in situ hybridization (**Figure S10A-F**). The podocyte-specific *Col4a3* upregulation was reminiscent of the protective COL4A3 allele identified in a GWAS study of DKD (Salem et al., 2019). Pathways related to oxidative stress, mitochondrial function, lipid metabolism (PPAR signaling) and gluconeogenesis were enriched in diabetic mouse PCT (**Figure S11A**), with *Pck1* and *Gsta2* emerging as the top up-regulated genes in both mouse models, which we validated by immunofluorescence (**Figure S11D-E**) and in situ hybridization (**Figure S11F-H**). The significant number of differentially expressed genes in the PCT suggest these transcriptional changes may reflect early signs of disease that precede overt, histologically detecTable damage. The transcriptional changes were in general more similar between HFD and 10-week BTBR mice (rather than 5 weeks), with substantial changes in response to injury occurring within the BTBR model from 5 to 10 weeks (**Figure S9A-C**). Finally, outside the heavily studied parenchymal compartment, macrophages consistently exhibited transcriptional changes across mouse models, prioritizing them for further investigation.

### Kidney disease progression marked by the expansion of a Trem2-expressing macrophage population in obese mice

Macrophages are known to play crucial roles in both the progression and resolution of kidney disease (Wang and Harris, 2011). We found multiple macrophage subsets in the HFD and BTBR *ob*/*ob* mouse models, including ‘resident’ (Resident Mϕ 1-7) and ‘infiltrating’ populations (Zimmerman et al., 2019) (**Figure S12A-G**). We cross-referenced macrophage subsets between the two mouse models, using a multi-class random forest classifier (**Methods**) trained on the HFD model macrophages, and applied to cells from the BTBR *wt*/*wt* and BTBR *ob*/*ob* mice (**Figure 5A, Figure S12H**, **Methods**). Both mouse models had shared resident (Resident Mϕ-1, 2 and 5 matching 1:1, Mϕ-7 in BTBR matching HFD-model Resident Mϕ-3 and Mϕ-4) and infiltrating macrophage populations.

**Figure 5.**
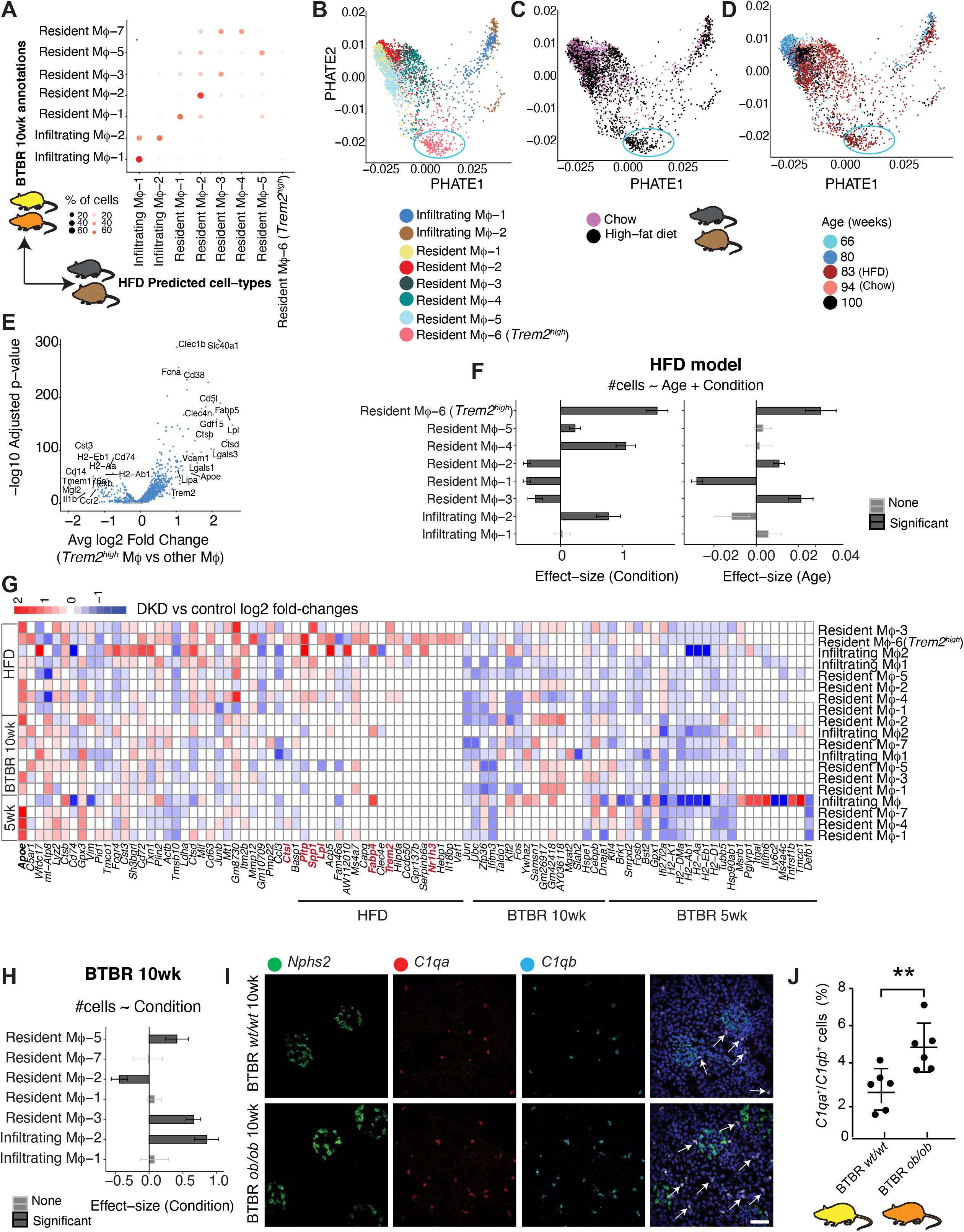
An increase in C1q-expressing macrophages in diabetic mice is associated with the expansion of a *Trem2-expressing* population in HFD-fed mice. (A) Comparison of macrophage populations between two mouse strains indicating shared and unique cell populations. Comparisons were performed by training a classifier on the HFD model macrophages and predicting labels on the 10-week-old BTBR macrophage data. (B-D) Potential of Heat diffusion for Affinity-based Transition Embedding (PHATE) visualization of macrophage populations in the kidneys of chow and HFD-fed mice. Macrophages are colored according to individual cluster (B), diet (C) and age (D). (E) Volcano plot showing DE genes in the *Trem2*^high^ population (resident Mϕ-6) in the kidneys of chow- and HFD-fed mice, compared to other kidney macrophages. (F-G) Effect-size of difference in macrophage subset proportions between conditions and ages in the (F) HFD and (G) BTBR 10wk mice. Standard error bars are shown. (H) *In situ* HCR using probes for *C1qa* (red) and *C1qb* (cyan) to identify resident macrophages in kidneys of 10-week-old BTBR *wt*/*wt* and *ob*/*ob* mice. The *Npsh2* probe green) was used to identify podocytes and provide spatial orientation. (I) Quantification of *C1qa+C1qb*+ expressing macrophages, expressed as a percentage of all cells, in kidneys of 10-week-old BTBR *wt*/*wt* (mean 2.7% +/- 0.4%, n=6) and BTBR *ob*/*ob* (mean 4.8% +/- 0.5%, n=6) mice. P values are derived from the unpaired Student’s t-test, **p<0.01. Arrows indicate individual cells. (J) Heatmap showing DE genes in resident and infiltrating kidney macrophage clusters in DKD (compared to control) across both mouse models (HFD and BTBR *ob*/*ob*) and time points in BTBR *ob*/*ob* mice (5 and 10 week). DE genes shared across all models are shown alongside model-specific differential expression. Color represents log2 fold changes of average gene expression in DKD over control in respective subsets.

Resident Mϕ-6 emerged as a unique population of macrophages expanded in obese, 90 week old HFD-fed diabetic mice (**Figure 5B-D**). This *Trem2^high^* population expressed *Cd5l*, a key mediator of lipid synthesis, and its transcriptional regulator *Nr1h3* (**Figure 5E, Figure S12C**), previously reported as a biomarker of DKD (Noelia et al., 2013; Peters et al., 2017). The resident Mϕ-6 cluster was also characterized by a signature that included *Trem2*, *Lpl*, *Fabp5*, *Lipa*, *Ctsb*, *Lgals3*, *Lgals1*, *Nceh1*, *Cd63* and *Cd36* (**Figure 5E**), also known as signature genes for lipid-associated macrophages (LAM) in the adipose tissue of HFD-fed, obese mice (Jaitin et al., 2019). *Trem2* and related genes were also differentially expressed identity markers in two additional kidney macrophage clusters in mouse kidney (resident Mϕ-4,-5), one of which had a signature associated with monocytic to LAM transition seen in adipose tissue (*Cd14*, *Ifrd1*, *C1qa*, *Cd63*, *Cd68*)(Jaitin et al., 2019). These *Trem2^high^* macrophage clusters, henceforth called “obesity-instructed macrophages,” expanded with disease progression (Poisson regression, **Figure 5F**, **Figure S12C**). A heat map of genes differentially expressed in disease across all mice (**Figure 5G**) highlighted the signature unique to the *Trem2*-expressing macrophage populations enriched for the PPAR signaling and lysosome pathways as has been implicated in adipose in the context of obesity. Subsets of resident macrophages were also expanded in 10-week BTBR *ob*/*ob* mice (**Figure 5H**, **Figure SI**), confirmed by *in situ* hybridization (mean *C1qa/b*+ macrophages 4.8% versus 2.7% in BTBR *wt*/*wt* mice, p<0.01; **Figure 5I-J**).

### An obesity-instructed TREM2^high^ macrophage population identified in kidneys of obese humans

We identified four subsets of macrophages (Mφ1-4) in combined data from human kidneys (**Figure 6A-B**). We then compared macrophage composition between mice and humans by using a multi-class random forest classifier (**Methods**) trained on the four subsets of human macrophages (**Figure 6A-B**) to classify mouse macrophages (**Figure 6C**). We found that over 85% of obesity-instructed *Trem2^high^* macrophage populations in mouse corresponded to human Mφ-4. Comparing the average gene expression profiles of the human *TREM2^high^* populations with the corresponding mouse HFD *Trem2^hĩgh^* populations (**Figure 6D**), showed high correlation (Spearman coefficient = 0.82). Species-specific divergence was noted for *APOC1* and *CD163* that were highly expressed in human Mφ-4, while *Lpl* and *Cd5l* were highly expressed in mouse *Trem2^high^* macrophages. Overall, this analysis revealed significant similarities in a specific obesity-instructed macrophage population shared between HFD-fed mice and humans.

**Figure 6.**
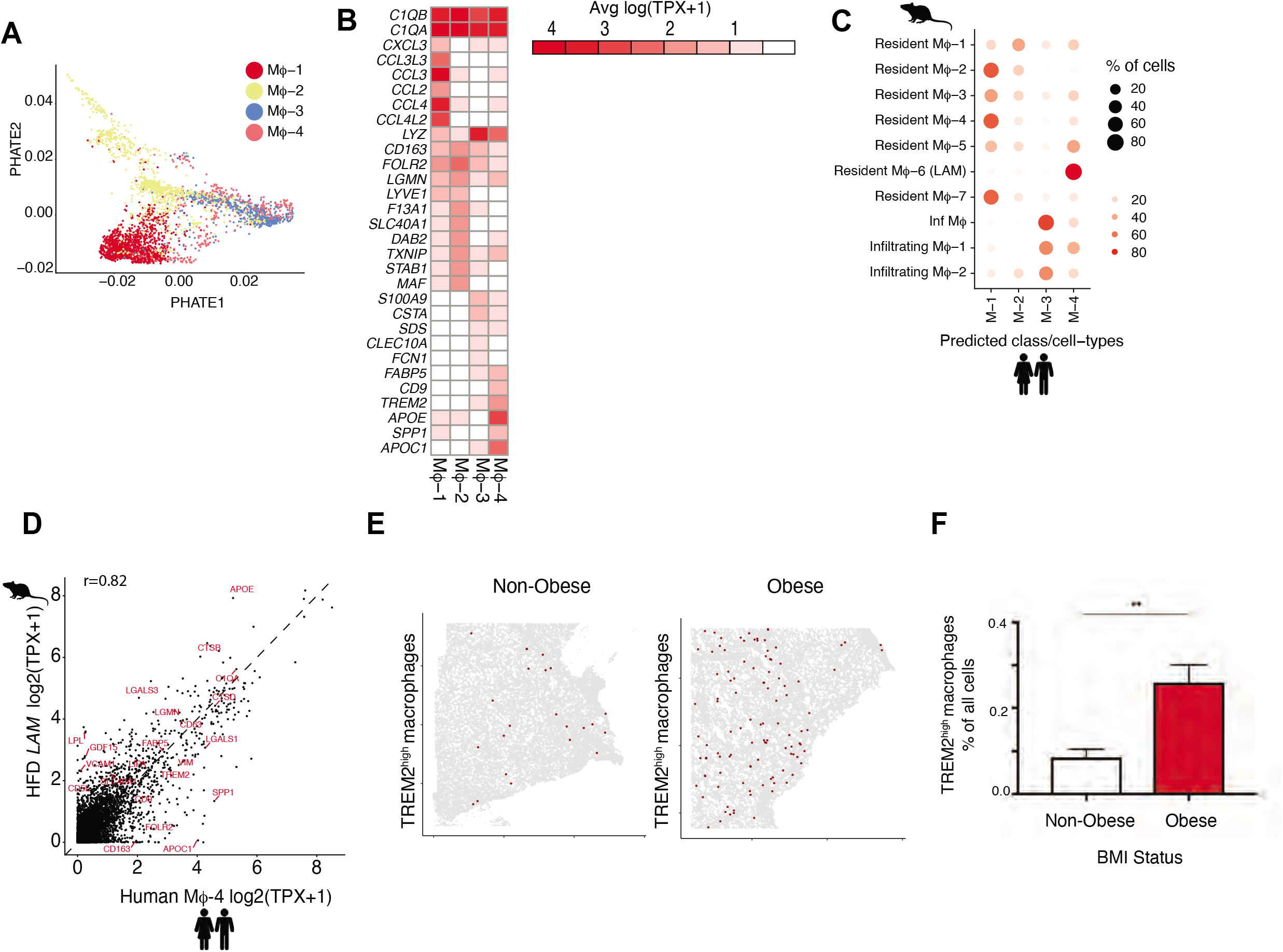
Expansion of a *TREM2^high^* macrophage population in kidneys of obese patients. (A) PHATE visualization of macrophages (MΦ) in the human diabetic kidney highlighting 4 populations (MΦ 1-4). (B) Heatmap visualization of expression programs distinguishing the 4 populations (MΦ 1-4). Rows are genes that are differentially expressed in MΦ 1-4. Values (color) represent row-normalized average gene expression in units of log(TPX+1). (C) Comparison of macrophage populations between patients and mouse strains to show shared and unique populations. Comparisons were performed by training a classifier on the human macrophages and predicting labels on the mouse macrophages. (D) Correlation of *TREM2*^high^ populations between HFD mouse (Y-axis) and human kidney (X-axis) shows shared and divergent genes. Each data point represents the average normalized and log-transformed expression of the gene in units of log(TPX+1). (E) Digital graphic representation of TREM2^high^ macrophages in nephrectomy tissue from non-obese and obese patients. Red circles denote TREM2^high^ macrophages with all cells (nuclei) shown in gray. (F) Quantification of TREM2^high^ macrophage populations in nephrectomy tissue from non-obese (mean 0.09 +/- 0.02%, n=5) and obese (mean 0.26 +/- 0.04%, n=6) patients. P values are derived from the unpaired Student’s t-test, **p<0.01.

To gain a deeper insight into Mφ-4, the macrophage population in humans that corresponded to *Trem2^high^* macrophages in mice, we sought to quantify TREM2 in human kidney tissue. TREM2^high^ macrophages were detected, dispersed throughout human kidney tissue sections (**Figure 6F**). Since these macrophages were uniquely detected in the HFD-fed, obese mouse model, we compared human kidney tissue from patients with high versus normal body mass index (BMI), the most widely accepted measure of obesity (Kato et al., 1996). This analysis revealed a significant increase in TREM2^high^ macrophages in kidney tissue from obese patients, as compared to kidney tissue from patients with a normal BMI (mean 0.09 +/-0.02% (n=5) v 0.26 +/-0.04% (n=^6^), p < 0.01; **Figure 6G**). Histologically, many of these patients had signs of kidney damage (as determined by an experienced renal pathologist), suggesting that this previously unknown TREM2^high^ macrophage expansion may coincide with the earliest signs of kidney injury. Our work has therefore revealed and molecularly defined a previously unrecognized, obesity-instructed macrophage population in human kidney, corresponding to an equally well-defined population in the HFD-fed mouse kidney.

## DISCUSSION

We have performed a high-resolution, cross-species comparison between mouse and human kidney that has revealed the shared transcriptomic architecture and cellular changes driven by obesity and diabetes. We specifically identified a population of obesity-instructed *TREM2^high^* macrophages in both human and mouse kidneys that track with kidney disease progression in the mouse. Our findings have significant implications for future therapeutic strategies, and lead to several conclusions.

First, we created a framework that allowed us to compare mouse and human kidney cellular taxonomies at high resolution, describing previously unreported granular subsets. This comparative atlas can serve as a resource for probing kidney transcriptional states across species. Overall, we concluded that broad cell class transcriptional identities are more highly conserved than granular cell subsets, especially podocytes, thick ascending limb cells and lymphatic endothelial cells. Transcription factor specificity scores suggested specific conservation in these cell types, whereas examples of divergence pointed to differential gene regulatory programs as the putative driver of inter-species differences. Hence, mechanistic investigations of mouse models must consider the granularity of the cell states for maximum translatability.

Second, we generated a comprehensive cell census (115,489 cells) of the obese and diabetic mouse kidney across two mouse models and time points, and compared disease signatures at the cell-type level. Again, while broad cell classes were conserved, each mouse model had its own granular subsets. Disease associated signatures were concordant between the HFD and BTBR kidney (at 10 weeks of age) in the cells of the nephron, with the distal tubular signature also overlapping with the human counterpart. Resident macrophages expanded in disease across both mouse models, but HFD-fed mice were superior as a model system in that they recapitulated the expansion of the *TREM2^high^* macrophage population that we also found in obese humans.

Importantly, our analysis distinguished specific populations of resident and infiltrating macrophages, including distinct populations of *LYVE1^high^* and *TREM2^high^* macrophages in both mouse and human kidney. The significance of resident macrophages, and especially *TREM2^high^* cells, in maintaining tissue homeostasis is increasingly being recognized (Cochain et al., 2018; Pinto et al., 2012). These cells have been shown to play a critical role in fending off inflammatory injury in adipose tissue, liver, and brain (Cochain et al., 2018; Jaitin et al., 2019; Keren-Shaul et al., 2017). Our identification of these cells in the kidney lends further support to the notion that this is an important, conserved population across different tissues, and across species.

We found that *TREM2^high^* macrophages had expanded in the kidneys of obese mice and humans. Recent work has revealed the TREM2 receptor as a major pathology-induced immune signaling node that senses tissue damage and activates robust immune remodeling as an adaptive response to injury (Deczkowska et al., 2020). Trem2-deficient mice have increased susceptibility to inflammation, obesity, and neurodegenerative disease (Deczkowska et al., 2020; Hammond et al., 2019; Jaitin et al., 2019). Our findings therefore suggest that these *Trem2^high^*, obesity-instructed macrophages in the kidney may be important determinants of the local response to kidney injury.

Emerging experimental and epidemiological evidence implicates obesity as an independent risk factor for kidney disease (Henegar et al., 2001). A recent study drawing from 10,547 patients showed that elevated BMI (over 25 kg/m^2^) accelerates the progression of kidney complications in patients with type 2 diabetes (ClinicalTrials.gov; NCT00145925) (Mohammedi et al., 2018). A limited number of studies have prospectively examined the relationship between BMI and adverse kidney events among individuals who have already developed type 2 diabetes (Belhatem et al., 2015; Huang et al., 2014; Luk et al., 2008). Furthermore, bariatric surgery improves kidney function (Chang et al., 2016; Hou et al., 2013; Navarro-Díaz et al., 2006). Our findings encourage follow up studies focused on preserving and promoting TREM2^high^ macrophages in order to protect kidney function in patients with both obesity and diabetes.

In sum, our cross-species comparison of mouse and human kidney cells provides a putative target (TREM2) and a model system (HFD-fed mice) for proof-of-concept studies with direct translatability to human obesity-driven kidney disease. More broadly, our work highlights how we can harness single cell technologies to zero in on novel targets for diseases with significant unmet medical need, and enhance the translatability of our findings from mouse models to patients.

## Acknowledgments

We thank our colleagues in the BWH Department of Urology for their generous assistance with recovery of tissue from tumor nephrectomies under IRB protocol 2011P002692. This work was funded by NIH grants DK095045 and DK099465 (AG), the Chan Zuckerberg Foundation (CZF019-002447) (AG), and the Slim Initiative for Genomic Medicine in the Americas (SIGMA) (AG).

## Author contributions

Conceptualization: YZ, AS, KV, AR, AG. Methodology: JLM, MA, FZ, MS, JW, MSS, DD, LTN, MSC, DD, JP, KK, SHS, EG, AG, AW, ACV, SLC. Software and Formal Analysis: AS Investigation: YZ, AS, KV, FZ, JLM Project administration: AS, KV, YZ, JLS, AG, AR Supervision: AG, AR. Writing: AG, AR, AS, JLS, KV, YZ

## Data and materials availability

Processed sequencing (scRNA-Seq) data will be deposited in the GEO database (https://www.ncbi.nlm.nih.gov/geo/), the single-cell portal for interactive visualization, and raw human sequencing data will be deposited in the controlled access repository DUOS (https://duos.broadinstitute.org/), upon publication.

## Materials and Methods

### Animals

#### Baseline mice

C57BL/6J mice were purchased from the Jackson Laboratory (JAX stock #000664; Bar Harbor, USA) and housed at 25°C with a 12-hour light-dark cycle in the American Association for Laboratory Animal Care (AALAC)-approved animal facility at the Broad Institute.

#### HFD Model

C57BL/6J mice (JAX stock #000664) were purchased from Jackson Laboratory (Bar Harbor, USA) and backcrossed with 129S1 mice (JAX stock #002448) for at least three generations, resulting in a C57BL/6J-129 mixed background strain. All mice were housed at 25°C with a 12-hour light-dark cycle in an AALAC-approved animal facility at Brigham and Women’s Hospital, Harvard Medical School. For the high-fat diet-induced DKD mouse model, adult C57BL/6J-129 mice were fed with a standard chow or high-fat diet *ad libitum*, starting at 8-10 weeks of age. The high-fat diet was purchased from Envigo (TD.93075 dough), which contains 55% per Kcal fat (23% saturated, 32% trans, 30% monounsaturated, 12% polyunsaturated), and 9.6% sucrose. To confirm the response and diabetic status of the HFD-fed mice, body weights were measured every two weeks after the commencement of the HFD. The intraperitoneal glucose tolerance test (IPGTT) was performed at 30 weeks of age and the intraperitoneal insulin tolerance test (IPITT) one week later. The 24-hour urine collection and scRNA-seq experiments were performed at 70, 80, 90, and 100 weeks of age.

#### BTBR model

3- and 6-week-old BTBR *wt*/*wt* and *ob*/*ob* mice were purchased from the Jackson Laboratory and housed as above at Brigham and Women’s Hospital, Harvard Medical School or the Broad Institute. The intraperitoneal glucose tolerance (IPGTT) and intraperitoneal insulin tolerance (IPITT) tests were performed in both BTBR *wt*/*wt* and *ob*/*ob* mice at 4 and 8 weeks of age (along with cholesterol levels) (Figures S4K-M), approximately 1 week prior to scRNA-seq experiments (Figure 4A). The 24-hour urine collection was performed at 5 and 10 weeks of age at the time of scRNA-seq.

All animal experiments were performed in accordance with the guidelines established and approved by the Animal Care and Use Committee at the Broad Institute (Animal Protocol No. 0061-07-15-1) and Brigham and Women’s Hospital, Harvard Medical School (Animal Protocol No. 01538).

### Glucose tolerance test

Prior to the test, mice were fasted for 16 h and transferred to a procedure room midway through the light phase of the light-dark cycle. Blood was obtained from a tail cut and assessed for baseline glucose levels using a One-touch UltraMini (Lifescan) glucometer. The mice then received 1 g/kg body weight of a 100 mg/ml glucose solution (Sigma, #G8769) in sterile PBS, delivered by i.p. injection. At 15, 30, 60, 90, and 120 min after the administration of glucose, dried blood and tissue were quickly removed from the tail wound and blood was collected again for glucose quantification.

### Insulin tolerance test

Prior to the test, mice were fasted for 5 h and transferred to a procedure room midway through the light phase of the light-dark cycle. The mice received 1 U/kg body weight of humulin N solution (Eli Lily #HI-310) in sterile PBS, delivered by i.p. injection. Blood glucose levels were assessed in the same way as GTT, before and 15, 30, 60, 90, and 120 min after the administration of humulin.

### Serum parameters measurement

Whole blood was collected in a 1.5 mL centrifuge tube. After collection, the whole blood was allowed to clot by leaving it undisturbed at room temperature for 15 min. The sample was then centrifuged at 1,000 x g at 4°C for 10 minutes. Following centrifugation, the supernatant was immediately transferred into a clean polypropylene tube on ice. If the serum was not analyzed immediately, it was divided into 100 μl aliquots and stored at –80°C. The levels of insulin, adiponectin, leptin, total cholesterol, and triglycerides were measured by the Animal Metabolic Physiological Core at Beth Israel Deaconess Medical Center. The levels of BUN and creatinine were measured by the Biochemical Genetics and Metabolic Disease Laboratory at the University of Alabama (Birmingham).

### 24-hour urine collection and urine albumin assay

Mice were housed individually in a metabolic cage supplied with adequate food and water. Urine was collected into a 25 mL falcon tube over 24 h. Total urine volume was measured and then centrifuged at 3,200 x g for 10 min at 4°C. Albumin quantification was done according to our previously published protocol (Zhou et al. 2017). Coumassie Blue stained gels of urine samples were quantified by densitometry with albumin standards using ImageJ software.

### Body Composition by Dual-Energy X-ray Absorptiometry (DEXA)

The mice were anesthetized using VetFlo™ Isoflurane Vaporizer (Kent Scientific Corp.). Their body composition was assessed using the PIXImus2 X-ray unit (GE Lunar Corporation, Madison, WI) connected to a computer equipped with LUNAR PIXImus2 software. The head region of each mouse was excluded from the analysis. The following parameters were automatically measured or calculated by the PIXImus2 software: bone mineral density (BMD), bone mineral content (BMC), bone area (B Area), tissue area (T Area), percentage fat, and total tissue mass (TTM). Fat mass, the percentage of lean mass, and lean mass, were calculated manually as follows: % Lean = 100 - % Fat; Lean Mass = TTM x (100 - % Fat); Fat Mass = TTM x % Fat.

### Light microscopy

Kidney tissue was fixed in 4% PFA overnight, embedded in paraffin and sectioned at 5 μm. Samples were deparaffinized and hydrated to water. Sections were oxidized in 0.5% periodic acid solution (Sigma-Aldrich) for 5 minutes, rinsed with distilled water and then placed in Schiff reagent (Sigma-Aldrich) for 15 min. They were then washed in tap water for 5 min and counterstained with Mayer’s hematoxylin (Sigma-Aldrich) for 1 min. Subsequent washing with tap water for 5 minutes was followed by dehydration and coverslip mounting using a synthetic mounting medium.

### Immunostaining

6 μm kidney sections were prepared using a cryostat. Samples were fixed with 4% PFA at room temperature and permeabilized with 0.1% PBS-Triton X for 10 min and blocked with 3% BSA at room temperature for 1 h. Samples were then incubated with primary antibodies (1:250 dilution for rabbit anti-Pck1 (# 720266, ThermoFisher Scientific, Waltham, USA) and rabbit anti-Gsta2 (# PA5-96757, ThermoFisher Scientific)) at room temperature for 1 h and washed 3 times with 0.1% PBS-Tween 20 for 10 min. Appropriate Alexa secondary antibodies (1:250 dilution for goat antirabbit IgG 594, # A32740, ThermoFisher Scientific) were used to visualize the proteins. Images were taken using an Olympus FV-1000 confocal microscope (Olympus America Inc, Center Valley, USA) and relative signal intensities were quantified using ImageJ software.

### Electron microscopy

Mouse kidney samples were prefixed with 4% PFA and then washed with PBS, prior to/following 2.5% glutaraldehyde (in PBS, pH 7.4) fixation at 4°C overnight. After washing in cacodylate buffer, kidney fragments were then postfixed in 1% osmium tetroxide for 1 h, dehydrated through ascending grades of alcohol, and embedded in Epon resin (Electron Microscopy Science, Hatfield, PA). Ultrathin sections (60 to 100 nm) were cut on an EM UC7 ultramicrotome (Leica Microsystems, Mannheim, Germany), stained with uranyl acetate and lead citrate, and examined with TEM (Morgagni 268D, Philips, Brno, Czech Republic).

### HCR and tissue collection

PBS perfused kidneys from wild-type mice of a 129/C57BL/6J hybrid background, BTBR *wt*/*wt*, and BTBR *ob*/*ob* mice were covered in OCT frozen in liquid nitrogen cooled isopentane. Thin sections of tissue (10um) were mounted in 24 well glass bottom plates (82050-898, VWR) coated with a 1:50 dilution of APTES (440140, Sigma). All HCR v3 reagents (probes, hairpins, and buffers) were purchased from Molecular Technologies. The following solutions were added to the tissue: 10% formalin (100503-120, VWR) for 15min, 2 washes of 1x PBS (AM9625, ThermoFisher Scientific), ice cold 70% EtOH at −20°C 2 hours to overnight, 3 washes 5x SSCT (15557044, ThermoFisher Scientific with 0.2% Tween-20), Hybridization buffer (Molecular Technologies) for 10min, probes in Hybridization buffer overnight, 4 15min washes in Wash buffer (Molecular Technologies), 3 washes 5x SSCT, Amplification buffer (Molecular Technologies) for 10min, heat denatured hairpins in Amplification buffer overnight, 3 15min washes in 5x SSCT (1:10,000 DAPI TCA2412-5MG, VWR in the second wash), and storage/imaging in 5x SSCT. Imaging was performed on a spinning disk confocal (Yokogawa W1 on Nikon Eclipse Ti) operating NIS-elements AR software. Image analysis and processing was performed on ImageJ Fiji.

### Imaging and analysis of mRNA expression amplified by *in situ* HCR

All data presented was acquired from multiple, distributed fields of view within each of the probed samples using either a Nikon Plan Fluor 40x 0.75 NA (*Col4a3*, *Col4a2*) or a Nikon Plan Apo 10x 0.45 NA Objective (*Gsta1/2*, *Pck1*). At each region of interest, 15 z-series optical sections were acquired with NIS Elements AR 4.51 software (Nikon, Tokyo, Japan) using a step size of 0.5 microns. The gamma, brightness, and contrast for all fluorescence micrographs were adjusted in NIS Elements software identically for compared sets of images.

Image analysis was performed on the maximum z-projections of the images acquired using custom MATLAB scripts (see https://github.com/ssturner-broad/HCRImageAnalysis.git) and the Image Processing Toolbox (MathWorks, Natick, USA). Briefly, the fluorescence signal corresponding to a gene of interest was first isolated and the background was subtracted. Fluorescence intensity was quantified as the integrated mean intensity of pixel values in a given region of interest. To compare fluorescence intensity quantitatively between experimental conditions, the intensity values for each field of view were normalized by the estimated number of nuclei identified in z-projections of paired DAPI channels. Cell density estimations were performed by detecting nuclei using an optimized algorithm for image segmentation and connected components detection. Finally, the resultant relative fluorescence intensity values were uniformly scaled across the compared sets of imaging data and expressed in terms of arbitrary units. For each of the genes visualized, measurements were based on no fewer than three independent fields of view for each sample. To evaluate statistical significance between the conditions compared in image sets, unpaired student’s t-tests were performed for each of the gene sets presented.

### Fluorescence image acquisition and analysis

All fluorescence imaging on human kidney tissue was performed using the Opera Phenix High-Content Screening System (PerkinElmer). 5mm cryosections were placed into individual wells of 24-well plates and the entire specimen initially imaged for DAPI at 5X, using the PreciScanTM feature (Perkin Elmer) to identify tissue. Pre-identified tissue regions were then imaged at higher resolution (20X water immersion objectives, confocal mode). Image analysis was performed using Harmony software (PerkinElmer).

### Single cell isolation from mouse kidney

Mice were anesthetized using isoflurane and perfused with PBS (1x) via the left ventricle. The kidneys were then removed and immediately placed in PBS on ice. For coronal sectioning, the renal capsule was removed, the kidney mounted and 300μm-thick sections cut using a vibrating blade microtome (Leica Biosystems Inc., Buffalo Grove, USA). The coronal sections were then returned to ice-cold PBS ready for dissection. With regards the regional sectioning, the renal capsule was removed from the other kidney and the hilum sampled first (figure 1B). The kidney was then cut in half along its coronal axis and cortical and medullary tissue was sequentially sampled. As each region was sampled, the tissue was chopped into 1mm x1mm cubes and placed in 0.25mg/ml liberase TH (Roche Diagnostics, Indianapolis, USA) dissociation medium. Following further dissection, the samples were incubated at 37°C for 1 hour at 600rpm. Samples were regularly triturated during the incubation period using a 1ml pipette, after which 10% heat-inactivated FBS RPMI was added to stop the digestion. Centrifugation at 500 g for 5 minutes at room temperature with the removal of the supernatant, was followed by the addition of ACK lysing buffer to remove erythrocytes (Thermo Fisher Scientific, Waltham, USA). Following centrifugation, the resulting cell pellet was incubated with Accumax at 37°C for 3 minutes (Innovative Cell Technologies Inc, San Diego, USA). 10% FBS RPMI was again used to neutralize the Accumax and centrifugation was followed by resuspension of the cell pellet in 0.4% BSA/PBS. The single cell suspension was filtered using a 30um CellTrics filter (Sysmex America Inc, Lincolnshire, USA) with cell viability and concentration determined using trypan blue and the Auto T4 cellometer (Nexcelom Bioscience LLC, Lawrence, USA). According to the manufacturer’s guidelines, 17,500 cells were loaded into the 10x Genomics microfluidic system/onto the 10x Genomics platform to achieve a targeted recovery of 10,000 cells (10x Genomics, Pleasanton, USA).

### Single cell isolation from human kidney tissue

Samples of macroscopically normal cortex, medulla and hilum were obtained from tumor nephrectomy specimens, distant from the tumor site and after appropriate patient consent, in accordance with IRB and institutional guidelines. Following transfer in 2% FBS RPMI, tissue was cut into 1mm x 1mm cubes and placed in 0.25mg/ml liberase TH (Roche Diagnostics, Indianapolis, USA) dissociation medium. Following further dissection, the tissue was incubated at 37°C for 1 hour at 600rpm as described previously. Samples were again regularly triturated during the incubation period using a 1ml pipette, after which the digestion was stopped by the addition of 10% heat-inactivated FBS RPMI. The addition of ACK lysing buffer (Thermo Fisher Scientific, Waltham, USA) following centrifugation at 500g for 5 minutes at room temperature, was performed twice in light of the lack of perfusion prior to nephrectomy. After centrifugation, the cell pellet was incubated with Accumax at 37°C for 3 minutes (Innovative Cell Technologies Inc, San Diego, USA), with 10% FBS RPMI again used for its subsequent neutralization. The resulting cell pellet was resuspended in 0.4% BSA/PBS and filtered using a 30um CellTrics filter (Sysmex America Inc, Lincolnshire, USA). Cell viability and concentration were determined as before, with 10,000 cells loaded into the 10x Genomics microfluidic system according to the manufacturer’s guidelines (10x Genomics, Pleasanton, USA).

### Droplet-based sn/scRNA-seq

Single nuclei or cells were partitioned into gel bead-in-emulsions (GEMs) and incubated to generate barcoded cDNA by reverse transcription. Barcoded cDNA was then amplified by PCR prior to library construction. Fragmentation, sample index and adaptor ligation, and PCR were used to generate libraries of paired-end constructs according to the manufacturer’s recommendations (10x Genomics, Pleasanton, USA). Libraries were pooled and sequenced using the Illumina HiSeq X system (San Diego, USA). Whenever feasible, we pooled 10x libraries on sequencing lanes to ensure that any individual sample was not confounded by batch (kidney section, day of sample collection, condition, timepoint) and were randomly distributed across lanes.

### Computational Methods for Data Analysis

#### Study Design and DKD cell atlases

##### (i) Mouse Baseline Kidney Atlas

For establishing the baseline atlas, the most commonly used lab mouse strain C57BL/6J between ages of 10 and 12 weeks, was used. We included biological replicates (n=5 for coronal section, n = 3 for renal hilum, medulla, cortex) from both male (M) and female (F) sexes. A mouse either donated a coronal section or a section each of cortex, medulla and hilum.

##### (ii) DKD Mouse Study Design

For the BTBR ob/ob genetic mouse model, we included n=5 and n=3 biological replicates for the 5- and 10-week time-point respectively for both *ob*/*ob* and wildtype strains. For the High-Fat diet (HFD) model, we combined n=4 biological replicates into a pooled “aged” group, in each of the chow and HFD conditions. The “aged” group ranged from 66 to 100 weeks old (**schematic Figure 4A**). For 3 biological replicates in each condition (Chow: ages 94,100; HFD: ages 83,100), there were 2 technical replicates or 10x channels run from the same cortical sample leading to a total of 7 libraries per condition.

##### (iii) Human kidney samples

Human kidney samples arrived as per the nephrectomy surgery schedule, with no selection on the age or sex of the patients. Where possible, the pathologist collected cortical, medullary and hilar sections (Supplementary Table 1). In case of 3 patients (Nx12, Nx10 and Nx8, the kidney sections were split to allow CD45 enrichment of one half. Though libraries were prepared soon after sample collection, we randomized pooling of samples during sequencing.

##### (iv) Sequencing Design

We pooled the 10x libraries on sequencing lanes using a randomized design to ensure that replicates from an individual batch (mouse, replicate, kidney section, day of sample collection, condition, timepoint) were distributed across lanes. Despite a deliberate random sequencing design to mitigate batch effects, donor effects were prominent in human data. Within a mouse strain, batch effects (separation by mouse) was not observed.

#### Generation of mouse and human baseline cell atlases

(*Panels in figures 1, Supplementary Figures 1-6*)

##### Preprocessing of 10x droplet based sequencing outputs

The mouse baseline, human baseline, HFD/Chow mouse, BTBR ob/ob mouse and human DKD scRNAseq data were generated via 10X Genomics Chromium 3’ droplet-based sequencing. The *Cellranger* toolkit (v2.1.1)(Genomics, n.d.) by 10X Genomics was used to (1) de-multiplex the sequencing outputs using the command *cellranger mkfastq* with default options, (2) align the sequencing reads to the reference transcriptomes (mm10 for mouse, GrCh38 for human), and (3) quantify gene expression, resulting in gene-by-cell UMI count matrices. (2) and (3) were done using the command *cellranger count* with default options.

##### Quality Control (QC) of scRNAseq datasets

Across all datasets, as an initial QC, we only retained cells that had reads mapping to a minimum of 1000 UMIs and 200 genes.

Because the kidney has the second highest mitochondrial metabolic activity among organs in the human body (Mercer et al. 2011), we first imposed a permissive threshold, allowing all cells with less than 80% mitochondrial reads to be included in the downstream analysis. We plotted boxplots showing the number of UMIs, genes and percent mitochondrial content after these filters.

We hypothesized that each cell type will have its own profile of metabolic activity and filtering must proceed in a cell type specific manner, leveraging exploratory data analysis. We proceeded to perform downstream steps and cluster cells (see **Inferring cell-types from mouse and human kidney cells** section).

First, we examined if there were any clusters that could be distinguished by a majority of cells expressing high-percentages of mitochondrial reads, and by examining the top differentially expressed genes for each cluster. In every (baseline mouse, btbr 4 week, btbr 10 week, HFD mouse models, human kidney data) model, we did not observe such a “technical” cluster. Because of the absence of technical clusters, we proceeded to assign high-level cell-type assignments to the clusters. On plotting percent mitochondrial reads across cells, we saw cells with higher percentages in cell types with high expected metabolic activity like the proximal tubule, thick ascending limb and the distal convoluted tubule. For any downstream analysis involving sub clustering or assessment of differentially expressed genes in disease, we discarded cells with over 20% mitochondrial reads except the proximal tubules which is explained in more detail below.

While ambient RNA did not pose a challenge for cell type annotations, we found ambient contamination during differential expression analysis in the mouse data. Specifically, genes highly expressed in the predominant parenchymal cell types proximal tubule and distal nephron, and some endothelial genes were frequently found to be ambient, i.e., differentially expressed in non-native cell types. We generated a list of such genes (“Pck1”, “Pecam1”, “Gsta2”, “Klk1”, “Spink1”, “Wfdc2”, “Miox”, “Slc34a1”, “Kap”, “Aldob”, “Cubn”, “Ttc36”, “Umod”, “Fxyd2”) and flagged them during differential expression analysis except in the native cell types where they are expected.

##### Inferring cell-types from mouse and human kidney cells

###### Normalization, Scaling and Dimensionality reduction

All single cell data analysis was performed in the R statistical computing environment (v3.5) using package *Seurat* (v2.3) (Butler et al. 2018). First, we normalized the gene UMI counts per cell using total sum scaling followed by multiplication by a factor of 1e+5 (TP10k or TPX, here we will refer to TPX), and log-transformation using a pseudocount of 1 using the *NormalizeData* function to obtain log(TPX+1) units for each gene. Next, we identified high varying features of gene expression using the *FindVariableGenes* function using default parameters. We computed the top 100 principal components (PCs) using the *RunPCA* function on the expression matrix composed of only the most highly variable genes after mean centering and scaling. We were more permissive in the number of PCs to enable discovery of rare cell types and states, as it is easier to *post hoc* merge together redundant states.

###### Unsupervised determination of putative cell types or clusters

For identification of cell types, we ran unsupervised clustering using the *FindClusters* function on all the computed PCs. In general, we used domain knowledge and exploratory data analyses to aid choice of parameters for the unsupervised analysis (resolution of clustering, annotation of cell types and number of PCs) *FindClusters* builds a shared k-nearest neighbor graph (Jarvis and Patrick 1973)(k=30) followed by Louvain community detection(Blondel et al. 2008) to determine clusters. We first used a resolution of 0.4, followed by iterative clustering on major cell types and as needed. For visualization, we embedded cells in the Uniform manifold approximation and projection (UMAP) space using the *RunUMAP* function using all computed PCs, and with default parameters.

###### Joint analyses of human cells from multiple donors

Batch effects are often detected by segregation of clusters by technical (e.g. donor origin, replicate, day of sequencing etc) rather than expected biological identity. While there were no explicit clusters from a technical replicate in the mice data, the human data separated by donor. We used Seurat’s *MultiCCA* function setting the number of canonical components to 20, and using variable genes present in at least 2 human individuals, to co-embed human scRNAseq data prior to clustering and visualization. Each individual was used as a separate batch.

###### Assignment of cell-identity

We ran differential gene expression (DGE) tests using the *FindAllMarkers* function to compute highly differentially expressed (DE) genes distinguishing each cluster from all other cells using the Wilcoxon Rank-Sum test with False Discovery Rate (FDR) adjustment using the method of Benjamini-Hochberg (Benjamani and Hochberg 1995). We performed the test only on genes expressed in at least 25% of the cells using the *min.pct* argument. For the joint analysis of the baseline mouse kidney, the *FindAllMarkers* function could not scale to the data size, and we used the base R *wilcox.test* function to compute DE genes, including only genes expressed in at least 25% of the cells using custom code. We annotated clusters by checking the presence of literature derived cell-type specific genes among the top DE genes when possible. Where applicable, Seurat’s *AddModuleScore* and *CellCycleScoring* functions were used to score signatures of gene sets and cell-cycle genes respectively.

###### Iterative clustering of mouse and human baseline atlas endothelial, immune and nephron epithelial cells

Iterative clustering was performed on endothelial cells (defined as cells in clusters where majority of the cells express *Pecam1/PECAM1*), immune cells (defined as cells in clusters where majority of the cells express *Ptprc/PTPRC*), mesenchymal cells (defined by a cluster where majority of the cells express *Myl9/MYL9*) and larger kidney parenchymal cells (proximal tubular, distal tubular, loop of Henle and collecting duct), by first subsetting cells in each category and then “subclustering” them. On sub-clustering, we checked that cluster membership was not exclusive to a single replicate. Sub-clustering was performed using the *FindClusters* function as described at a resolution of 1. Often, this distinguished expected higher-level myeloid and lymphoid, and endothelial classes (fenestrated vs not, progenitor-like etc). In case of over clustering of subsets, we merged cell types that had a single gene as DE to the next higher level of cell classes in order. One round of sub clustering was often sufficient but in case of rare cells or certain expected subclasses (lymphoid) or when a subset of a cluster was distinct (e.g. neutrophils, human pDCs) on visualization, an additional round of sub clustering on specific clusters was performed.

###### Flagging of ectopic cells

In the mouse hilum, we found cells expressing the cholesterol and lipid metabolism genes *Cyp11a1, Cyp11b1, Cyp21a1, Star* and *Akr1cl*, typical of the adrenal gland, and hence assigned them as “Ectopic Adrenal,” consistent with reports of ectopic adrenal tissue being found under the capsule in rodent kidneys (Frazier et al. 2012). We recovered a set of *Aqp7*+ endothelial cells exclusive to the renal hilum, reported to be associated with glycerol transport (Skowronski et al. 2007); however, because of their absence in the coronal section, this set may also be ectopic unless further verified (Supplementary Figure S2B and D). Ectopic cells were not included in downstream comparisons.

###### Identification of rare cells

To identify the juxtaglomerular apparatus (JGA) cells, we performed iterative clustering on mesenchymal cells in the mouse cortex, using *Ren1* as a handle. By iteratively clustering the cortical mesenchymal cells, we identified three major subsets: *Ptn* and *Itga8*-expressing mesangial cells which we localized to the mouse glomerulus using HCR (**Figure S4C**), vSMCs expressing *Acta2* and *Myl9*, and pericytes expressing *Rgs5*. The pericytes could be further divided into Reninexpressing (*Ren1+/Ren^hi^*) and Ren1-/Ren^low^ cells. For identifying the *macula densa*, we used *Nos1/NOS1* expression as a handle to decide which cluster to sub cluster among *Slc12a1/SLC12A1*+ cells.

###### Curation of gene signatures

We used literature derived genes for kidney parenchymal, stromal and immune cell types (Stewart et al., 2019; Subramanian et al., 2019; Young et al., 2018).

###### Identification of cell type markers

We derived cluster specific DE genes using the *FindAllMarkers* function to distinguish cell-types at the higher and subcluster levels (Figure S2A-D). For visualization on the heatmap, we selected the both data-driven markers obtained by DGE (at least top 3 in case of immune cells) in addition to literature-derived canonical markers.

###### Plotting and visualization

- We used the *ggplot2* (Wickham 2016) R package for generation of boxplots (*geom_box*), violin plots (*geom_vln*), proportional bar plots (*geom_col*), UMAP visualization and dotplots (*geom _point*).
- In the dot plot representation, each dot size represented the percentage of cells expressing the gene, and the color represented the average nonzero gene expression in log2 scale.
- Cell proportions are computed as the ratio of the number of cells in a certain cell type divided by the total number of cells in the unit of interest (e.g. condition, region, individual, Figures G,L,Q, Supp Figure.1P-R).
- We visualized heatmaps using the *pheatmap* (Kolde 2012)package. Units of expression are log transformed transcript counts per 10,0000 (TPX). A pseudocount of 1 was added during the log transformation.

#### Characterization of macrophages in human adult kidney *(Panels in figures 2, Supplementary Figures 3-4)*

We iteratively clustered C1Q+ positive macrophages to derive subsets, followed by DGE analysis using the *FindAllMarkers* functions to determine subset specific signatures.

##### Receptor-ligand analysis

Putative ligands for LYVE1 and TREM2 were selected from the Fantom database, CellphoneDB database and literature. Only those cell subsets where either receptor or ligand was present in at least 25% of the cells were determined to have robust expression. Dotplots were plotted as described above.

##### Cross-tissue analysis of macrophages

We downloaded publicly available (Microwellseq) human adipose, liver, heart, and kidney scRNAseq datasets, as post processed “gene x cell” expression raw count matrices from NCBI GEO (GSE134355, (Han et al. 2020)). For each tissue, we separately performed normalization, HVG selection, scaling, dimensionality reduction and clustering as described above. We then subsetted the macrophages as both CD45+ and C1QA/C1QB+ cells. Human microglia expression matrices (MARS-Seq) were downloaded from NCBI GEO GSE134707 (Geirsdottir et al. 2020). We combined the normalized gene expression matrices of the macrophages across the 5 public datasets along with the human adult kidney macrophages, performed HVG selection, scaling, dimensionality reduction, co-embedding using the *RunHarmony* function of the *Harmony* (Korsunsky et al. 2019) R package using the technology (10x, MARS-Seq or Microwellseq) as batch, graph-based clustering on the top 20 Harmony dimensions using the *FindNeighbors* and *FindClusters* function (resolution=0.5) of *Seurat* v3 (Stuart et al. 2019), and DGE of the clusters as described above. We removed contaminating clusters of B-cells, fibroblasts, endothelial and epithelial cells. We subclustered the remaining cells and used UMAP to visualize the resulting five clusters, and barplots to represent the composition of each of the 6 clusters by tissue, and type of human kidney macrophage encoded as the human kidney macrophage subsets, and “others” for other cells.

#### Comparison of mouse and human kidney in health *(Panels in figures 3, Supplementary Figure 7)*

##### Derivation of orthologous genes

The database of mouse and human orthologous genes was downloaded from the Mouse Genome Informatics (Smith and Eppig 2012) website (To Check: citation http://www.informatics.jax.org/downloads/reports/index.html#homology). We note here that there are limitations to using orthologous genes, as genes of interest may be missing or synonym gene names may be used. We added one obvious edit: *Ctgf:CTGF*, which appeared as its synonym *Ccn2*, whereas our dataset had the gene listed as *Ctgf*. Another point to note is that in case of paralogs, only one gene is listed and this may represent one of several possible orthologs; of note *Gsta2* is one such gene which had *GSTA5* listed as its human ortholog.

##### Random-Forest classifier based correspondence of cell types between mouse and human kidney

We trained a multi-class random forest classifier (RF) on the baseline mouse sagittal data, using the cell-type labels as the classes, using the *randomForest* function in the *randomForest* R package, as previously demonstrated in (Shekhar et al. 2016; Peng et al. 2019). We applied the classification task on three datasets: (1) Cd45-/Cd31-, (2) Cd31+, and (3) Cd45+. In each case, we split the mouse dataset into training and test sets. The training set was composed of a random sample from each cell subset, with the sample size being the minimum of either 70% of the cells or 1000. The remaining held out data in each class composed the test set. For features, we chose the intersection of highly variable orthologous genes from both the baseline human and mouse datasets. The gene expression matrices in units of log(TPX+1) were scaled to mean 1 and variance 0 prior to training and testing. The minimum sample size was set as the smallest training set class, and we chose the option of 5000 trees with the function call *“randomForest(x=training_data, y=training_label, importance = TRUE, ntree = 5001, proximity=TRUE, sampsize=sampsizes, keep.inbag=TRUE, replace=FALSE*)”. The trained RF was then used to predict the mouse cell type label for each human cell. The confusion matrix of the resulting predictions were visualized as a dotplot, where the size and color of the dots referred to the proportion of cells in the test dataset cell subset (rows) assigned a given mouse cell subset label (columns). We considered a correspondence or mapping between a mouse and human cell subset to be (1) 1-1 if greater than 90% of the cells in the human subset mapped to the mouse subset, (2) orthologous and highly specific if 80-90% mapped (3) orthologous if 65-80% mapped and (4) non-specific if <65% mapped. We performed this analysis separately on the CD45-/PECAM1-, endothelial and immune compartments. We note that while a lack of specific mapping may reflect true non-specificity, it may also be reflective of “impure” clustering or inseparability either because of indiscrete classes or noise (ambient RNA or multiplets).

##### Assessment of transcription factor (TF) specificity

We downloaded the database of 1639 human TFs provided in (Lambert et al. 2018), and derived a subset of 1253 TFs having orthologs in mice, and in our data. We grouped the mouse and human cell types into broad cell class, and selected those categories present in both species (16 in CD45-/CD31-: Podocyte, PEC, PCT, tDL, TAL, DCT, CNT, CD-PC, CD-A-IC, CD-B-IC, Urothelium, Fibroblast, vSMC, Mesangial, Pericyte, Glia; 5 in CD31+: GEC, FEC, LEC, DVR, MGP+ELN+; CD45+: pDC, cDC, Macrophages, B Cells, Plasma Cell, T Cell) for further evaluation. We computed the % of cells in any subset that express the TFs (% expressed), and only selected those TFs that are expressed by at least 25% of the cells in any subset, leading to a smaller set of TFs: 380 among the CD45-/CD31-cells, 157 among CD31+ cells.

For each TF, the TFSS was defined as (1-SEI) where SEI is the Shannon equitability index. The Transcription Factor Specificity Score (TFSS) was computed as follows:

1. The percentage of cells expressing each TF was computed in each shared class within the compartment of interest.
2. the vector of % of cells expressed per TF was divided by the total sum of % expressed for normalization.
3. The Shannon diversity index was computed on the normalized % expressed vector.
4. (SEI) is computed as the ratio of Shannon diversity index divided by log2(total number of subsets).

We looked for TF activity divergence by selecting for TFs that had high specificity in one species but not in the other suggesting putative differential regulation. Assuming a linear relationship between TFSS of both species, divergent TFs were defined as those having a large residual from the linear regression fit using the command “resid(lm(mouse-human ~ 0))” in R where mouse and human represent the respective species TFSS.

#### DKD atlases in mouse and human (*Panels in figures 4*)

DKD atlases were generated following steps of dimensionality reduction by PCA, clustering and determination of differentially expressed genes as described for the baseline atlases. In each model -- HFD mice, BTBR mice at weeks 5 and 10 as well as human data -- both DKD and non-DKD replicates were combined during clustering. For the human data, donor effects were corrected using Seurat’s *MultiCCA* function as in the baseline atlas. In each mouse model, cells were partitioned into endothelial, immune, PCT and “other” cells first, then sub-clustered to identify granular subsets. In case of PCTs, in the BTBR models, a distinct cluster of cells with very high mitochondrial content and low number of genes expressed separated out, which we discarded.

#### Identification of DKD-associated genes and pathways *(Panels in Supplementary Figures 9-11, Figure 5G)*

##### Derivation of differentially expressed (DE) genes in DKD

In the mouse data, after assignment of cell-identity as described previously, we performed differential expression analyses between control and DKD cells using the Wilcoxon-Rank sum test using the “wilcox.test” base R function on genes present in at least 25% of cells in either condition, and a Benjamini-Hochberg FDR cutoff of 0.05 using the “p.adjust” base R function to prioritize for downstream interpretation. We did the analysis for each cell type independently. We analyzed five mice in each of the HFD- and chow-fed groups, pooled across 66-100 weeks of age into a single “aged mice” group for DGE analysis (**Figure 4A**).

For human data, we used a poisson mixed-effects model with the donor as the random covariate, disease as covariate being tested and the average number of UMIs per cell as the offset. We used the *glmer* command from the *lme4* package in R using the formula ‘gene~condition+offset(log(scalenUMI))+(1|nephrectomy).’ The model was run for each cell type independently. We adjusted for multiple hypotheses by the method of Benjamini and Hochberg. We prioritized genes by setting fdr < 0.1.

##### Pathway analysis

We ran *enrichR* on the prioritized DE genes to determine enriched gene sets using the KEGG database. We corrected for multiple hypothesis testing by the method of Benjamini and Hochberg (alpha < 0.05).

###### Comparison of DE genes in DKD across mouse models

(Figures 5G, Supplementary Figure 11A)

We first prioritized those DE genes with absolute fold change greater than 1 and adjusted p-value < 0.05, for each cell type. Fold-changes were computed as the log2 ratios of average normalized gene expression across all cells in a cell type. DE genes in some cell types did not pass these filters. To summarize results across the mouse models, we divided the prioritized differentially expressed genes in each cell type and model into 2 classes: (1) genes that belong to a cell type unique to a model; (2) genes that belong to a cell type shared between at least 2 models. We used a heatmap representation with rows as the selected genes, and columns as cell types to visualize the results, where each element was a fold-change.

#### Characterization of macrophages in mouse diabetic kidney *(Panels in figures 5, Supplementary Figures 12)*

##### Annotation of immune cells and macrophages

For both the HFD and BTBR mouse models (two time points), we subsetted the *Cd45*+ cells and iteratively clustered them to derive myeloid and lymphoid lineages. We used the previously characterized resident and infiltrating kidney macrophage signatures to mark macrophages as resident vs infiltrating (Figure) (Zimmerman et al. 2019). For the heatmap visualizations (Supplementary Figure 12 D-G), we used the top 3 data-driven DE genes marking the subset, alongside selected canonical and literature driven markers. On all heatmaps, gene expression in units of log(TPX+1) was averaged across all cells in a subset, followed by scaling across all subsets.

As macrophage subsets are in a transcriptional continuum, we used the Potential of Heat-diffusion for Affinity-based Trajectory Embedding (PHATE, (Moon et al. 2019)) embedding to visualize the cells across age, condition and subset (**Figure 5 B-D**). The macrophage genexcell data matrices were first converted to the anndata object format using the *anndata* Python library(https://anndata.readthedocs.io/en/stable/#), and PHATE was run using the *phate* Python library, and the commands *“phate.PHATE()”* and *“phate _op.fit_transform(anndata*).”

The Lipid associated macrophage (LAM) signature was derived from (Jaitin et al. 2019). We performed DGE using the Wilcoxon rank-sum test using the function call wilcox.test in R, between the HFD LAMSs and other HFD macrophages. The results were visualized as a volcano plot (**Figure 5E**), where genes that passed the FDR alpha threshold of 0.05 were colored blue, while the rest were gray. Top DE genes were annotated on the plot with gene names.

##### Comparison of macrophage populations across strains

We trained a RF classifier on the HFD macrophage subset labels using either the intersection of HVG with BTBR week 10 (Figure 5A) or week 5 (Supplementary Figure 12H) macrophages as features and then predicted on the BTBR data to determine correspondence.

##### Association of cell type proportions to age and disease

We used a Poisson regression model to determine the association between macrophage proportions (using counts) and covariates (age, disease condition). The model was defined as follows using the glm command from the stats package in R and formula “ncells~offset(log2(total_cells))+age+condition” for the HFD model and “ncells~offset(log2(total_cells))+condition” for the BTBR models.

#### Characterization of macrophages in human diabetic kidney (*Panels in figures 6*)

##### Derivation of human DKD macrophage subsets

We subsetted and clustered all the CD45+ immune cells across DKD and non-DKD kidney data, including the CD45+ enriched samples, and subsetted the clusters differentially enriched in the macrophage marker genes C1QA and C1QB into the human kidney macrophage subset. We then iteratively clustered the human kidney macrophage subset to derive 4 distinct subsets (MΦ 1-4), after excluding monocytes and DCs, and performed DGE using the FindAllMarkers Seurat command with the default Wilcoxon rank-sum test to get subset-specific marker genes.

We used PHATE to visualize the co-embedding of the cells, and a heatmap visualization of marker gene expression. For the heatmap, each marker gene was assigned the average gene expression across all cells in the subset in units of log(TPX+1), followed by row normalization across the subsets.

##### Comparison of mouse and human macrophages in DKD

To map the mouse macrophages with human macrophages, a multi-class RF was trained on the human DKD macrophages as described above on the feature set of orthologous HVG derived from the DKD mouse models and MΦ 1-4. The trained RF was used to predict labels for human macrophages. The resulting confusion matrix was visualized as a dot plot as described above.

To compare the *LYVE1*+ human MΦ-2 subset, with the *Lyve1*+ population in baseline mice, we examined the correlation of genes between the two subsets, by computing the Spearman correlation between the two average gene expression vectors. We repeated the correlation comparison between the HFD LAM-like macrophages and human MΦ-4. In both cases, gene expression in units of log (TPX+1) was averaged across the respective subsets in comparison, and plotted as a scatter (**Figure 3M, 6D**). Genes strongly deviating from the slope (species-specific) or lying on the slope (shared), were annotated.

##### TREM2 Graphic Images

TREM2 macrophages were selected as described previously. Representative regions were created by first finding the median x and y coordinates of all TREM2 macrophages in a section and setting a target-nuclei count of 40,000 total nuclei. The radius of the square, centered around the median that contained the target nuclei count, rounded to the nearest 10, was determined and represented in the graphic image. Graphs visualized using ggplot2.

