## Supplemental Data for "Obesity-instructed *TREM2^high^* macrophages identified by comparative analysis of diabetic mouse and human kidney at single cell resolution"

#### SUPPLEMENTARY FIGURES

##### Figure S1.

**Coronal sectioning captures an overview of the mouse kidney cellular taxonomy**
(A-E) UMAP visualization of *Cd45-/Cd31-* cells obtained via a joint embedding of the different kidney regions (A), cortex (B), medulla (C), renal hilum (D), and the coronal section (E). PEC, Parietal Epithelial cells; EGM, Extraglomerular Mesangial cells; PCT, Proximal Convoluted tubule; tDL, thin descending limb; tAL, thin ascending limb;TAL, thick ascending limb; DCT, Distal convoluted tubule; CNT, connecting tubule, CNT; Collecting duct cells: CD-PC, Principal; CD-A-IC, Intercalated alpha;CD-B-IC, Intercalated beta; IMCD, Inner Medullary Collecting Duct; vSMC, vascular smooth muscle cells.
(F-J) UMAP visualization of *Cd31+* endothelial cell classes obtained via a joint embedding of the different kidney regions (F), cortex (G), medulla (H), renal hilum (I), and the coronal section (J).
(K-O) UMAP visualization of *Cd45+* immune cells obtained via a joint embedding (K) of the different kidney regions, cortex (L), medulla (M), renal hilum (N), and the coronal section (O).
(P-R) Barplot representation of cell classes by region (P) *Cd45-/Cd31-*, (Q) endothelial and (R) immune cells.
On all UMAPs, points represent individual cells, colors represent cell subsets shown in legend.

##### Figure S2.

**Transcripts identifying *CD31-/CD45-* and *CD31+* endothelial cell subsets in mouse and human kidney with multiple PEC and PCT subsets**

(A, B) Heatmap visualization of the column-scaled average gene expression in log(TPX+1) units of canonical marker and data-derived differentially expressed genes specific to the *CD31-/CD45-* cell classes derived from the (A) human and (B) mouse regional kidney atlas.

(C, D) Heatmap visualization of the column-scaled average gene expression in log(TPX+1) units of marker genes defining endothelial cell subsets in the (C) human and (D) mouse kidney.

##### Figure S3.

**Transcripts identifying *CD45+* immune cell types in mouse and human kidney**
(A, B) Heatmap visualization of the column-scaled average gene expression in log(TPX+1) units, of marker genes identifying the immune cell subsets in the human (A) and perfused mouse (B) kidney.
(C) Dot plot representation of marker genes (columns) distinguishing the 5 macrophage human subsets. Size of the dots represent the percentage of cells expressing the gene (columns). Color of the dots represents the normalized average expression in units of log(TPX+1).
(D) UMAP visualization of resident MΦ pan-marker and subset specific marker genes in the kidney meta-atlas

##### Figure S4.

**Validation of newly characterized kidney macrophage and PEC populations**

(A) *In situ* HCR to demonstrate *LYVE1*$^{high}$ macrophages in human kidney, using probes for *LYVE1*, the homeostatic macrophage marker *FOLR2,* and PCT marker, *LRP2*, for spatial orientation.
(B) *In situ* HCR using probes for *TREM2* and *LGALS* to confirm the presence of *TREM2*$^{high}$ macrophages in human kidney tissue, with probes for *LRP2* providing spatial context.
(C-D) Violin plots of data-driven and canonical marker gene expression in units of log(TPX+1) distinguishing PEC (C) and PCT (D) subsets in mouse kidney: Parietal Epithelial Cells (PEC): *Cldn1*, *Tnc*, *Dkk3*, *Wt1*; Podocyte: *Nphs2*, *Nphs1*, *Wt1*; Proximal Convoluted Tubular (PCT) cells: *Lrp2*, *Cubn*, *Acsm3*, *Slco1a1*, *Slc22a8*.
(E, F) *In situ* HCR validation in C57BL/6J-129 control mice of scRNAseq-driven markers for PEC-1 *Dkk3* (E) and *Tnc* (F) alongside podocyte specific markers *Nphs2* and *Wt1*. (*Dkk3*, *Tnc*, cyan; *Nphs2*, green; *Wt1*, red).
(G, H, I) *In situ* HCR validation in C57BL/6J-129 control mice of scRNAseq-driven markers for PCT-S1 marker *Slc5a2* (G), PCT-S3 segment *Acsm3* (H) and *Slco1a1* (I) alongside pan-PCT marker *Lrp2* and the S1-S2 marker *Slc5a1* and *Slc22a8*. *Lrp2* also co-localizes with *Slc22a8*. (*Acsm3*, *Slco1a1*, *Slc5a2* cyan; *Lrp2*, green; *Slc22a8*, *Slc5a1* red).

**Figure S5.**

**PCT sexual dimorphism and enrichment of rare juxtaglomerular apparatus and mesangial cells**

(A) UMAP visualization of *Cd31-/Cd45*- cells in the cortical mouse kidney stratified by sex.
(B) Dot plot representation of marker genes (rows) distinguishing proximal tubular subsets (columns), highlighting those dominating male or female by symbols. Size of the dots represent the proportion of cells in each subset (columns) expressing the gene (rows). Color of the dots represents the normalized average expression in units of log(TPX+1).
(C) UMAP visualization of mouse kidney cortical mesenchymal cells. Sub-clustering resulted in four subsets, including juxtaglomerular apparatus (JGA) cells.
(D) Dot plot visualization of transcriptomic markers distinguishing the four mouse mesenchymal subsets. Size of the dots represent the proportion of cells in each subset (columns) expressing the gene (rows). Color of the dots represents the average gene expression in units of log(TPX+1).
(E) *In situ* HCR validation in C57BL/6J-129 control mice shows spatial expression of mesangial (*Ptn* and *Itga8*) and JGA (*Ren1*) markers juxtaposed with podocyte marker *Nphs2*.
(F, H) UMAP visualization of sub-clusters in the medullary distal nephron thick ascending limb (TAL) cells reveals a rare population of macula densa cells in mouse (F) and human (H).
(G, I) Dot plot visualization of transcriptomic markers distinguishing the macula densa of (G) mouse and (I) human kidney. Size of the dots represent the proportion of cells in each subset (columns) expressing the gene (rows). Color of the dots represents the level of average expression in units of log(TPX+1).
(J) *In situ* HCR validation in C57BL/6J-129 control mice of the *Nos1* macula densa marker juxtaposed with *Slc12a1*, marking the thick ascending limb (TAL).

**Figure S6.**

**Immune cell diversity in mouse kidney**

(A, B) Violin plots of data-driven marker gene expression in units of log(TPX+1) distinguishing resident macrophages from infiltrating macrophages and other immune cells. (A) *C1qa* (B) *C1qb*.
(C, D) *In situ* HCR in C57BL/6J-129 control mice was used to validate the presence of macrophages. Probes complementary to *C1qa* (C) and *C1qb* (D) were designed to identify resident macrophages, seen next to glomeruli and in the wider tubulointerstitium. *Nphs2* (podocin) and *Lrp2* (megalin) were used to provide concurrent localization of podocytes (i.e. glomeruli) and proximal convoluted tubules, respectively.
(E, H) Violin plots of data-driven marker gene expression in units of log(TPX+1) distinguishing pDCs and cDCs. (E) *Siglech* (pDC) (H) *Naaa* (cDC).
(F, G) *Siglech* expression (cyan) confirmed the presence of plasmacytoid DCs, seen alongside glomeruli (localized using *Nphs2,* green), lymphatic vessels (*Mmrn1*, red, top panel) and fenestrated endothelium (*Plvap*, red, bottom panel) using *in situ* HCR in C57BL/6J-129 control mice.
(I) Conventional DCs were localized using *in situ* HCR probes complementary to *Naaa* and seen in the tubulointerstitium with *C1qa*-expressing resident macrophages.
pDC, plasmacytoid dendritic cell; cDC, conventional dendritic cell; Res Mø, resident macrophage; Inf Mø, infiltrating macrophage; NKT, natural killer T cell.

**Figure S7.**

**Comparison of mouse and human broad cell classes and *in situ* validation of endothelial cells**

(A-C) Dot plots representing prediction results in human data of random-forest classifier trained on human CD45-/CD31- (A), CD31+ (B) and CD45+ (C) broad cell classes. The size and color of the dot represents the proportion of cells in a given mouse cell class (Y-axis) that was assigned a cell type label corresponding to or classified as the human cell class (X-axis).
(D, F) Violin plots of data-driven marker gene expression in units of log(TPX+1) distinguishing endothelial subsets in mouse and human kidney: PLVAP+ Fenestrated endothelial cells (FEC): Plvap/PLVAP; GEC: Ehd3/EHD3; lymphatic endothelial cells (LEC): Mmrn1/MMRN1, Lyve1/LYVE1.
(E, G) *In situ* HCR validation in C57BL/6J-129 control mice of (E) *Ehd3* (green), a marker of the glomerular endothelium, along with *Pecam1* (red), a global endothelial marker and *Plvap* (cyan), a marker for fenestration diaphragms; and (G) *Lyve1* (cyan) and another data-driven lymphatic endothelial marker *Mmrn1* (red).

**Figure S8.**

**Phenotypic characterization of mouse DKD models**

(A-J) Metabolic profile of chow- and HFD-fed mice: body weight (A), glucose tolerance test (B), insulin tolerance test (C), and plasma levels of insulin (D), adiponectin (E) and leptin (F). Body fat percentage (G), plasma cholesterol level (H), triglyceride level (I) and bone mineral density (J) are also shown. Chow (n = 33); HFD (n = 39); Mean +/- S.E.M. *$p<0.05$, ***$p<0.001$.
(K-L) Glucose (K) and insulin (L) tolerance tests in 5- and 10-week-old BTBR *wt/wt* and *ob/ob* mice, respectively. BTBR *wt/wt* 5wk n= 9; *ob/ob* 5wk n = 9; BTBR *wt/wt* 10wk n= 9; *ob/ob* 10wk n = 9; Mean +/- S.E.M.

(M-O) Measurement of renal parameters in 5- and 10-week-old BTBR *ob/ob* mice; 24h urinary albumin (M), blood urea nitrogen (BUN, N)) and serum creatinine (O). Mean +/- S.E.M. ***p<0.001
(P) Plasma total cholesterol levels in 5- and 10-week-old BTBR mice. Mean +/- S.E.M. ***p<0.001.
(Q) Transmission electron microscopy of kidney tissue from 10-week-old BTBR *wt/wt* and *ob/ob* mice.
(R) Assessment of podocytes in glomeruli from patients with and without DKD using *in situ* HCR, with probes complementary to podocin (*NPHS2*, red) and synaptopodin (*SYNPO*, green). *LRP2* (megalin, cyan) delineates the proximal convoluted tubules (DAPI, blue); scale bar 50µm. Podocytes were quantified in three independent fields of view for each of the DKD and non-DKD samples.
(S) Quantification of podocytes by *in situ* HCR in patients with and without DKD. Mean podocyte percentage 26.2% +/- 1.74% in non-DKD versus 18.1% +/- 0.27% in DKD, **p<0.01 (unpaired Student's t-test).

**Figure S9.**

**Differentially expressed genes for mouse models and human data**

(A-D) Number of differentially expressed genes (FDR < 5%) by cell class for the mouse models (Wilcoxon-rank-sum test): HFD (A), BTBR 5wk (B), BTBR 10wk (C) and the human data (D, poisson mixed-effects model, FDR<0.1).

(E-H) Upset plots representing shared and unique signatures among the human and mouse DKD data in the (E) DCT, (F) TAL, (G) podocyte and (H) mesangial cells. The set size represents the total number of differentially expressed genes in DKD. The intersection size represents the number of genes in the intersection of the sets represented by the black dots connected by the line segment.

**Figure S10.**

**Changes in podocyte and mesangial cell DKD-related gene expression**

(A) Boxplots showing DE of *Col4a3* in podocytes from the BTBR and HFD mouse models.
(B-C) Representative *in situ* HCR images (B) and quantification (C) of *Col4a3* expression in podocytes of 10-week-old BTBR *wt/wt* (n=3) and BTBR *ob/ob* (n=3) mice. Scale bars 50µm. **p < 0.01, unpaired Student's t-test. (*mean +/- SEM values to be added*)
(D) Boxplots showing DE of *Col4a2* in mesangial cells from the BTBR and HFD mouse models.
(E-F) Representative *in situ* HCR images (E) and quantification of *Col4a2* (F) in mesangial cells of 10-week-old BTBR *wt/wt* (n=5) and *ob/ob* (n=5) mice. Scale bars 50µm. ***p < 0.001, unpaired Student's t-test.

**Figure S11.**

**Shared and mouse model-specific transcriptomic changes in proximal convoluted tubular cells precede overt signs of injury by histology**

(A) Heatmap showing differentially expressed (DE) genes in the PCT cell subsets of kidneys from mice with DKD (compared to controls), in both mouse models (HFD and BTBR *ob/ob*) and time points (5- and 10-week-old BTBR mice). DE genes shared across all models are shown alongside model-specific differential expression. Color represents log2 fold changes of average gene expression in DKD over control in respective subsets.
(B) Heatmap representation of genes in human PCT cells orthologous to mouse DE genes (A) in the 3 patients with DKD (Nx#10, Nx#11, Nx#12). Color represents log2 fold changes of average gene expression in DKD over the average of the 9 patients without DKD.
(C) Kyoto Encyclopedia of Genes and Genomes (KEGG) pathways for which the DE genes in DKD (A) are enriched across all mouse models.
(D) Immunofluorescent staining of Pck1 (red) in kidney tissue from chow- and HFD-fed mice.
(E) Quantification of Pck1 immunofluorescence in kidneys from chow- (n=5) and HFD-fed (n=5) mice; $**p < 0.01$, FU, fluorescence units. Scale bars 50µm.
(F) *In situ* HCR for expression of *Pck1* (red), *Gsta1/2* (cyan) and *Lrp2* (green) in kidneys from 10-week-old BTBR *wt/wt* and BTBR *ob/ob* mice.
(G) Quantification of *Pck1* expression in 10-week-old BTBR *wt/wt* (n=4) and BTBR *ob/ob* (n=4) mice by *in situ* HCR; $*p < 0.05$, unpaired Student's t-test. RFU, relative fluorescence units.
(H) Quantification of *Gsta1/2* expression in 10-week-old BTBR *wt/wt* (n=4) and BTBR *ob/ob* (n=4) mice by *in situ* HCR; $***p < 0.001$, unpaired Student's t-test. RFU, relative fluorescence units

**Figure S12**
**Immune cell populations present in the kidneys of the HFD and BTBR mice**

(A-C) Proportions of immune cells in the kidneys of 5-week-old BTBR *wt/wt* and BTBR *ob/ob* mice (A), 10-week-old BTBR *wt/wt* and BTBR *ob/ob* mice (B) and chow- and HFD-fed mice (C). Individual populations are represented by distinct colors.
(D) Heatmap showing canonical and data-derived DE genes used for immune cell cluster annotation in chow- and HFD-fed mice.
(E) Heatmap showing canonical and data-derived DE genes used for 5-week-old BTBR *wt/wt* and BTBR *ob/ob* mouse kidney immune cell cluster annotation.
(F) Heatmap showing canonical and data-derived DE genes used for immune cell cluster annotation in 10-week-old BTBR *wt/wt* and BTBR *ob/ob* mice.
(G) Heatmap showing DE genes characterizing kidney resident and infiltrating macrophage populations in the two mouse models (HFD and BTBR) and time points (5- and 10-week-old BTBR).
(H) Comparison of macrophage populations between mouse strains to show shared and unique populations. Comparisons were performed by training a classifier on the HFD model macrophages and predicting labels on the 5-week-old BTBR macrophage data.
(I) Effect size of condition on macrophage subset proportions as determined by a Poisson regression model in the 5-week-old BTBR *ob/ob* mice.

**Supplementary Tables**

**Table S1:** Table of QC metrics, and metadata for mouse and human (in separate sheets for each species).

Clinical, histological and demographic details for non-DKD and DKD patients, including creatinine and proteinuria assessment were recorded.

**Table S2:** List of taxonomies for mouse and human by compartment, broad cell classes and granular subsets (in separate sheets for each species)

**Table S3:** List of differentially expressed genes for each granular subset in human

**Table S4:** List of differentially expressed genes for each granular subset in mouse

**Table S5:** List of differentially expressed genes for human macrophage subsets, along with breakup by nephrectomy.

**Table S6:** Shared broad cell classes

**Table S7:** DKD QCs + Shared broad cell classes

**Table S8:** DE genes in human DKD

**Table S9:** DE genes in HFD, BTBR wk 10

### Supplementary Figure 1

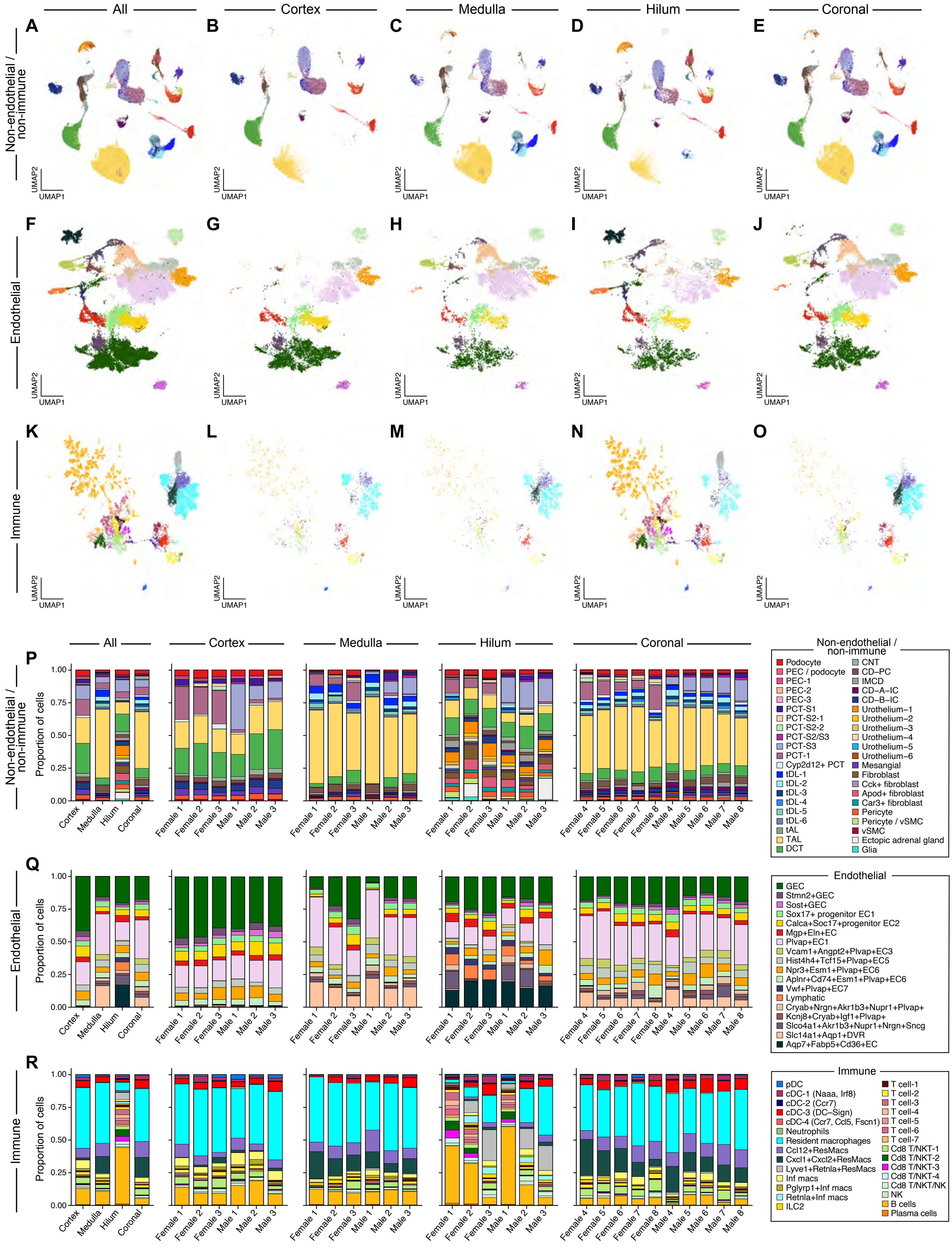

Supplementary Figure 2

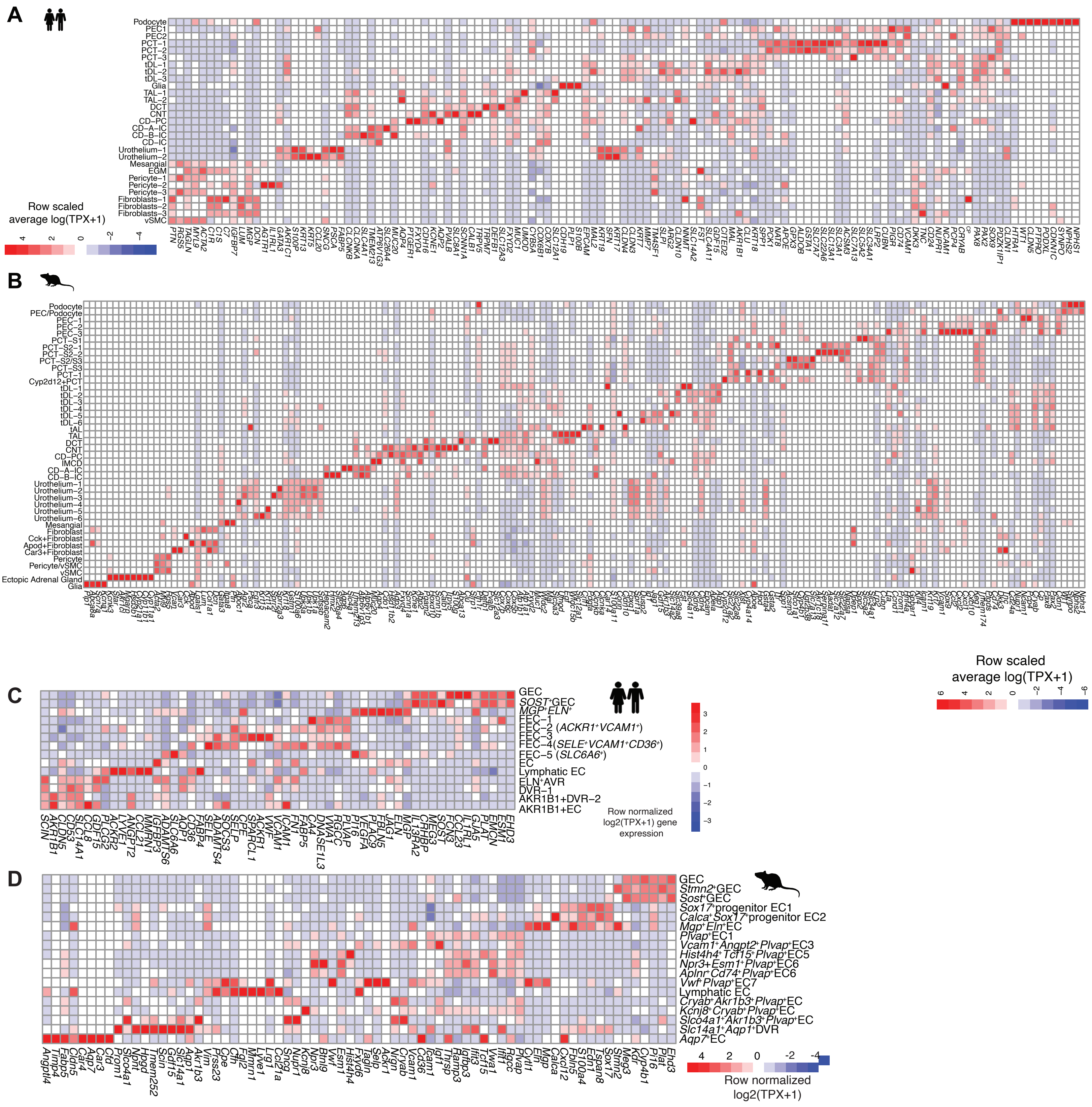

Supplementary Figure 3

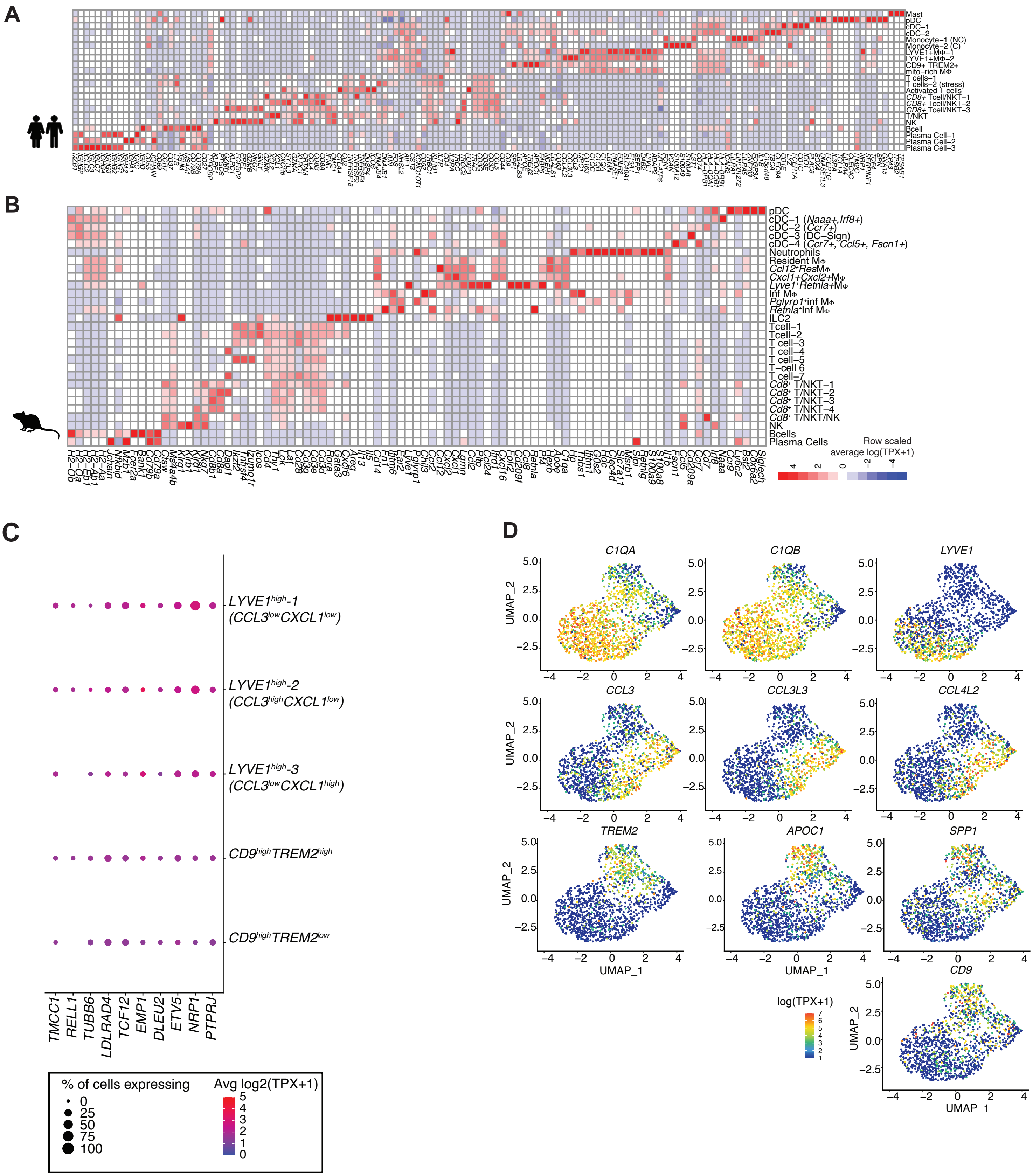

Supplementary Figure 4

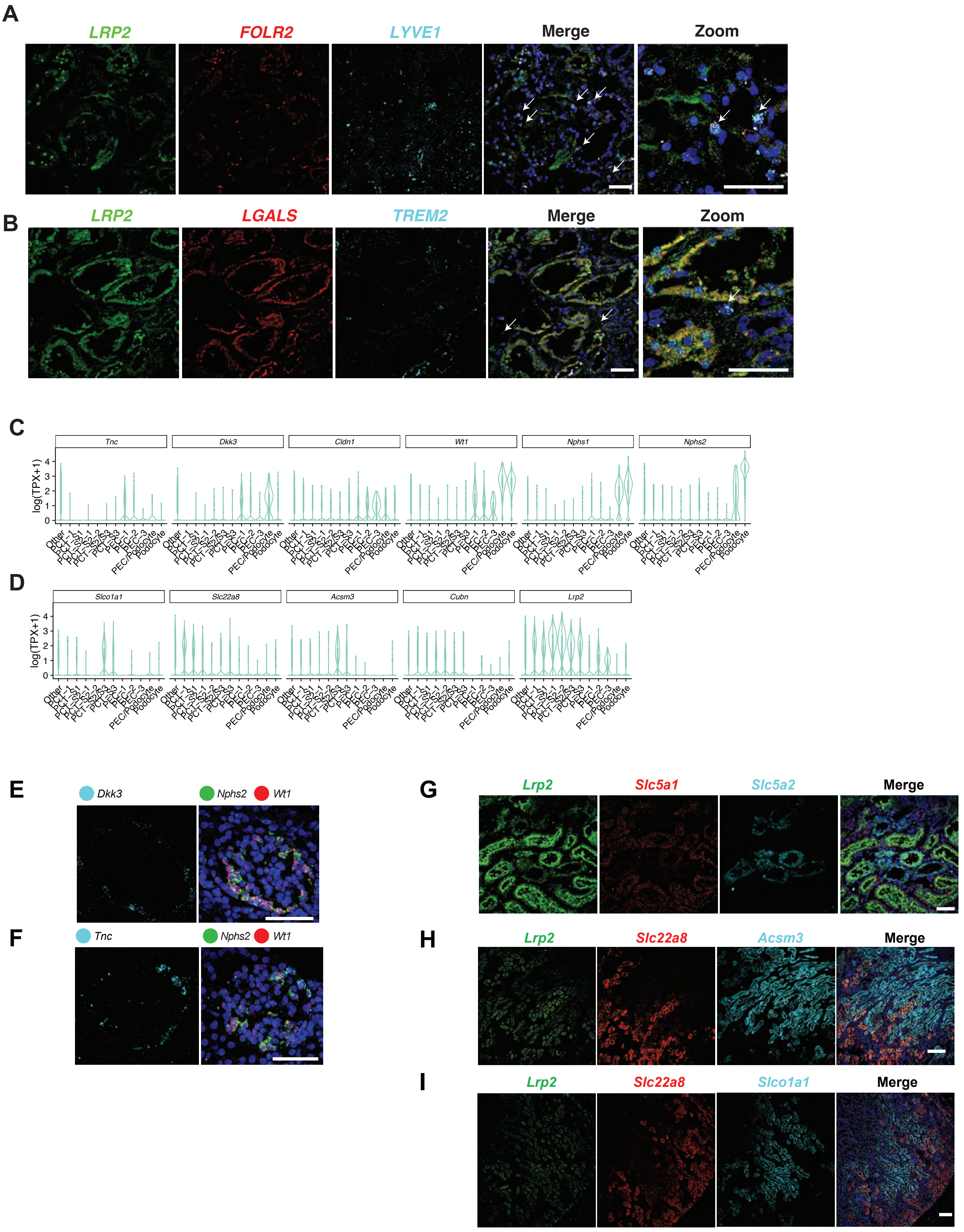

Supplementary Figure 5

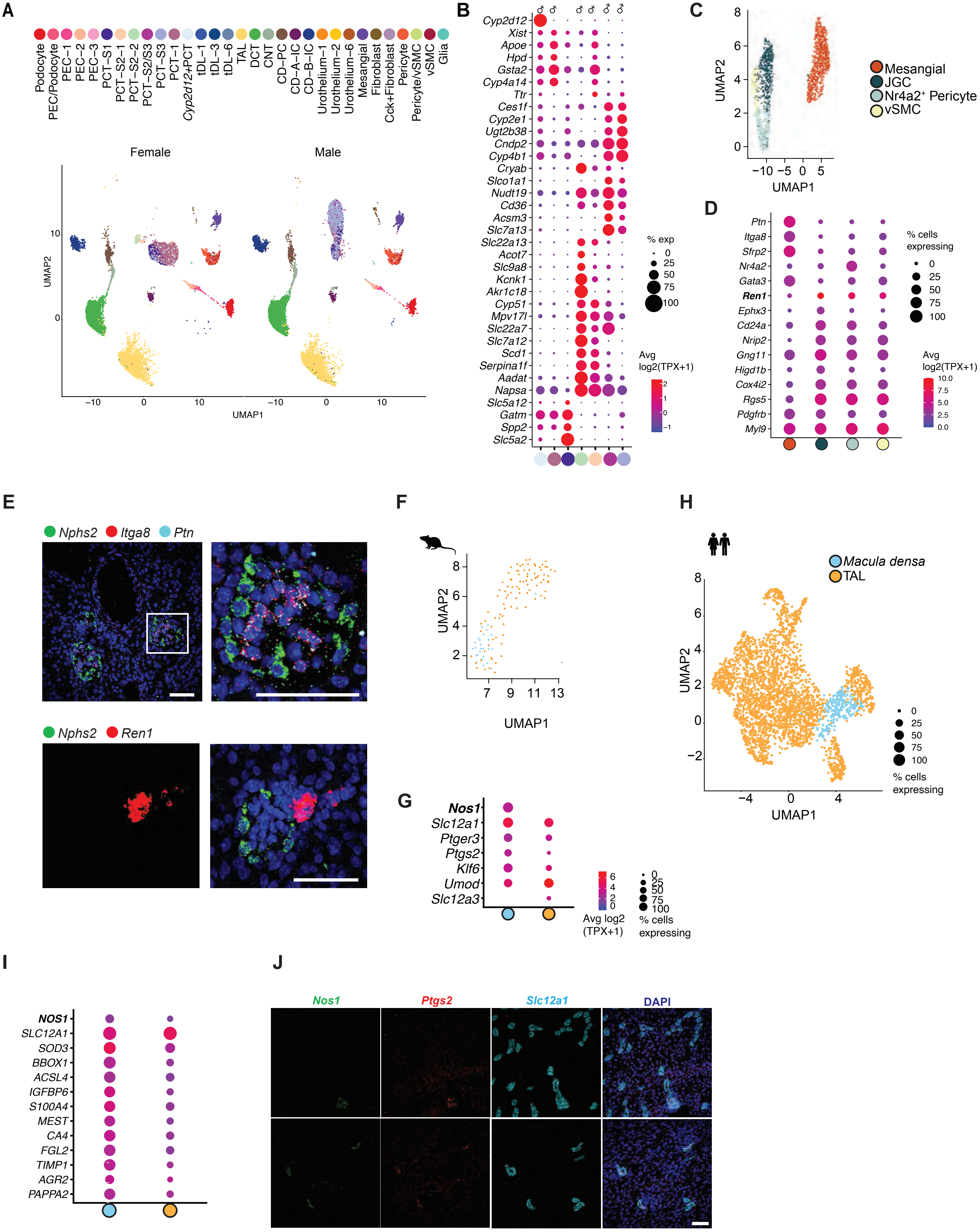

### Supplementary Figure 6

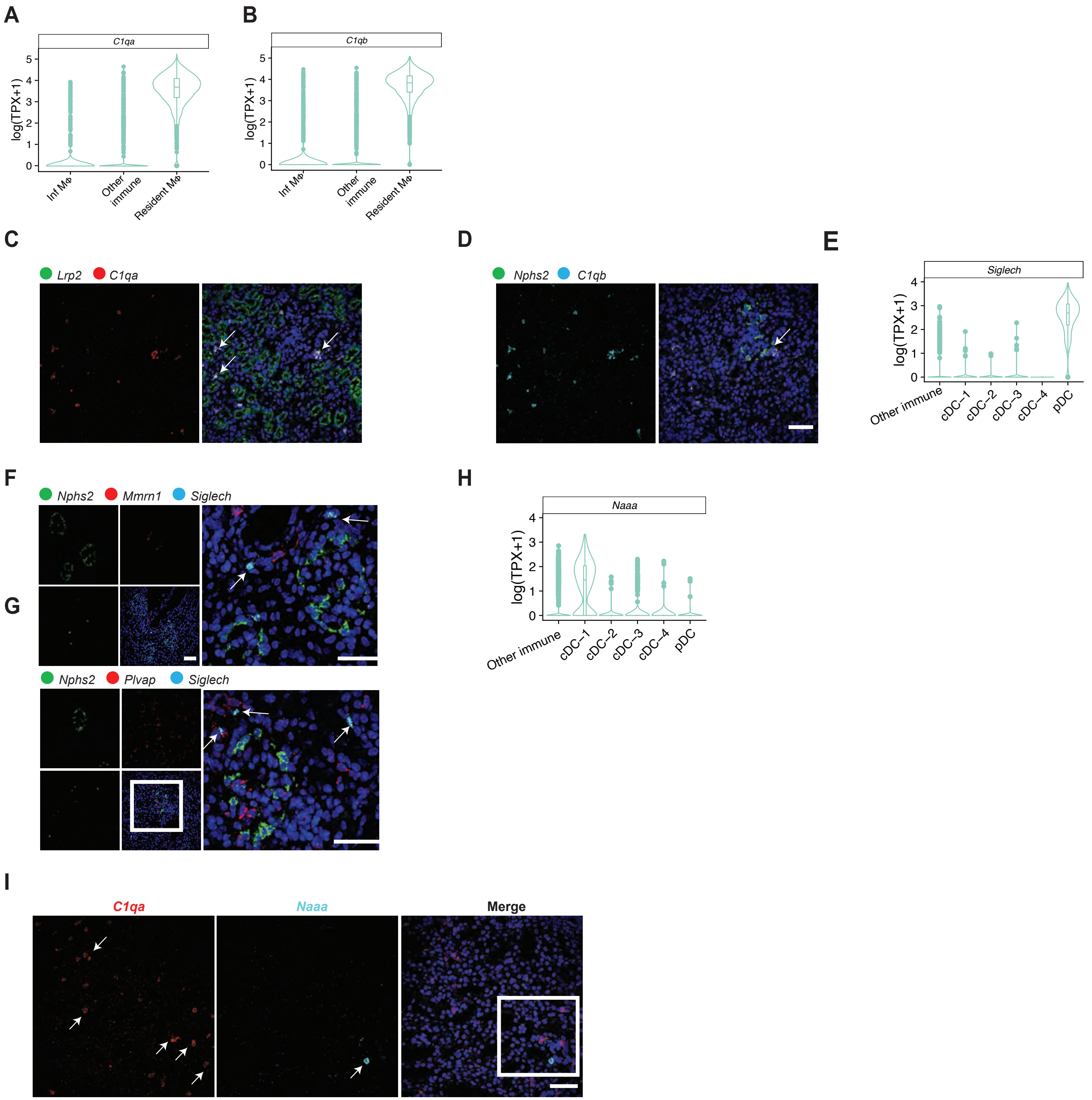

Supplementary Figure 7

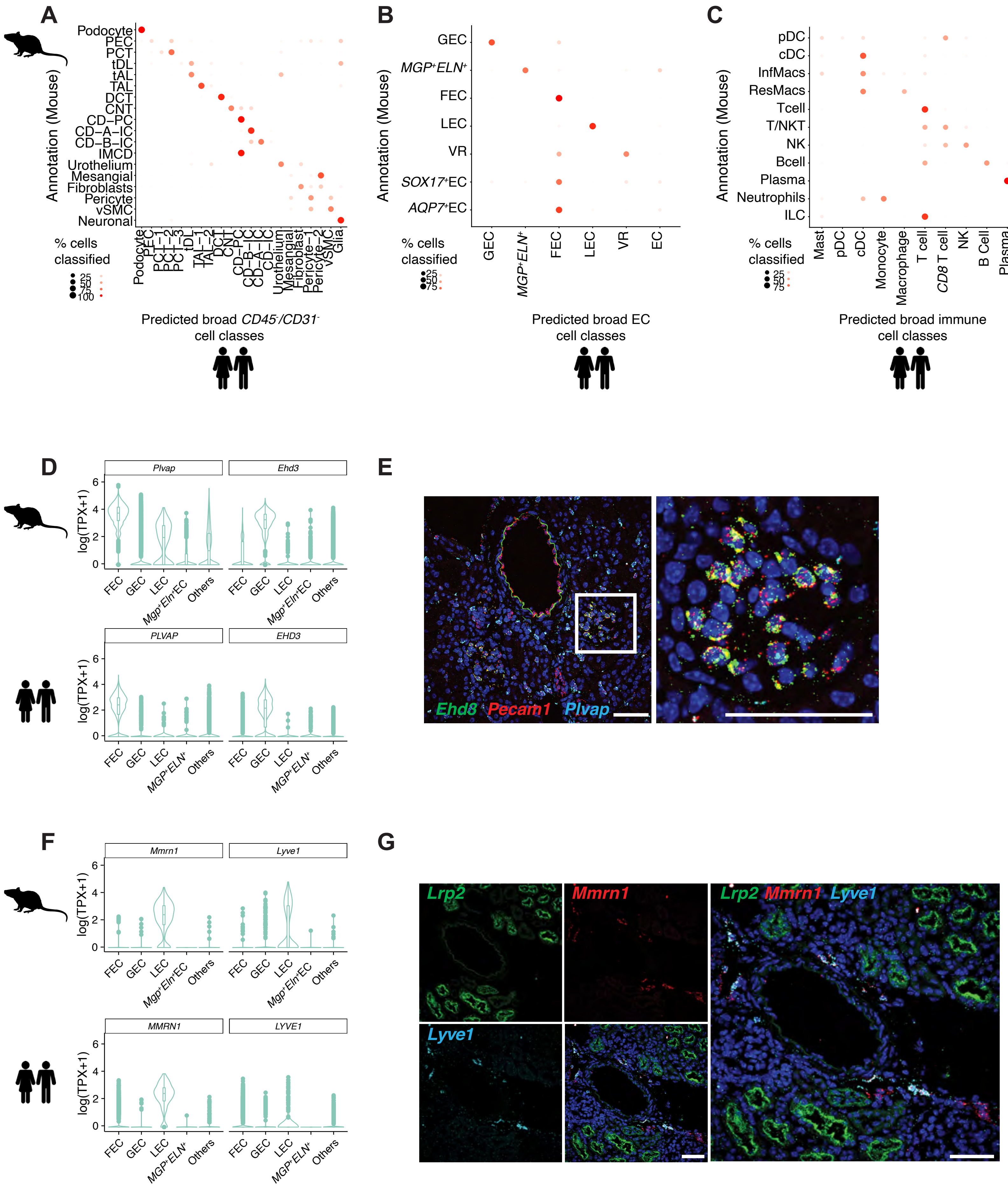

### Supplementary Figure 8

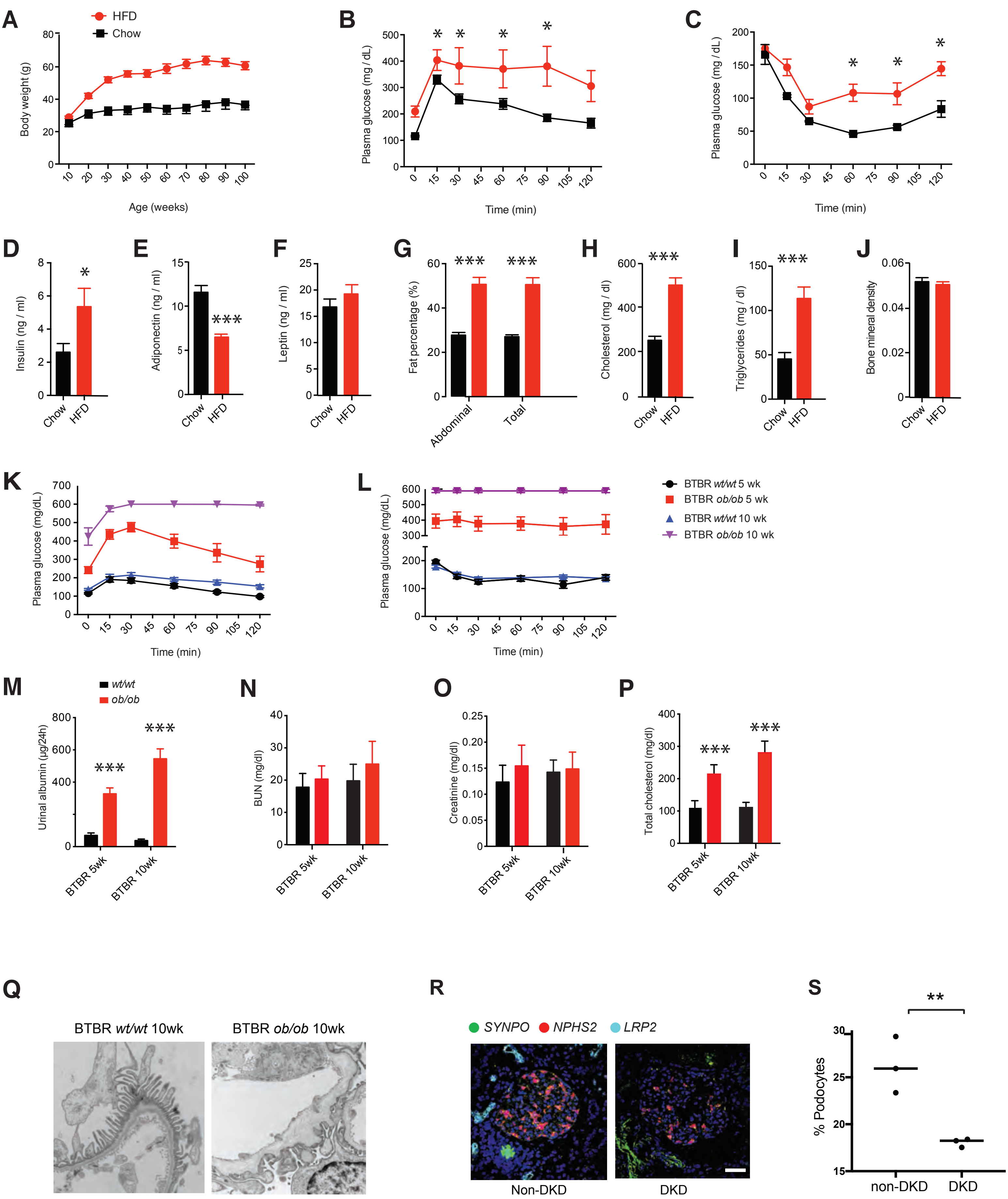

Supplementary Figure 9

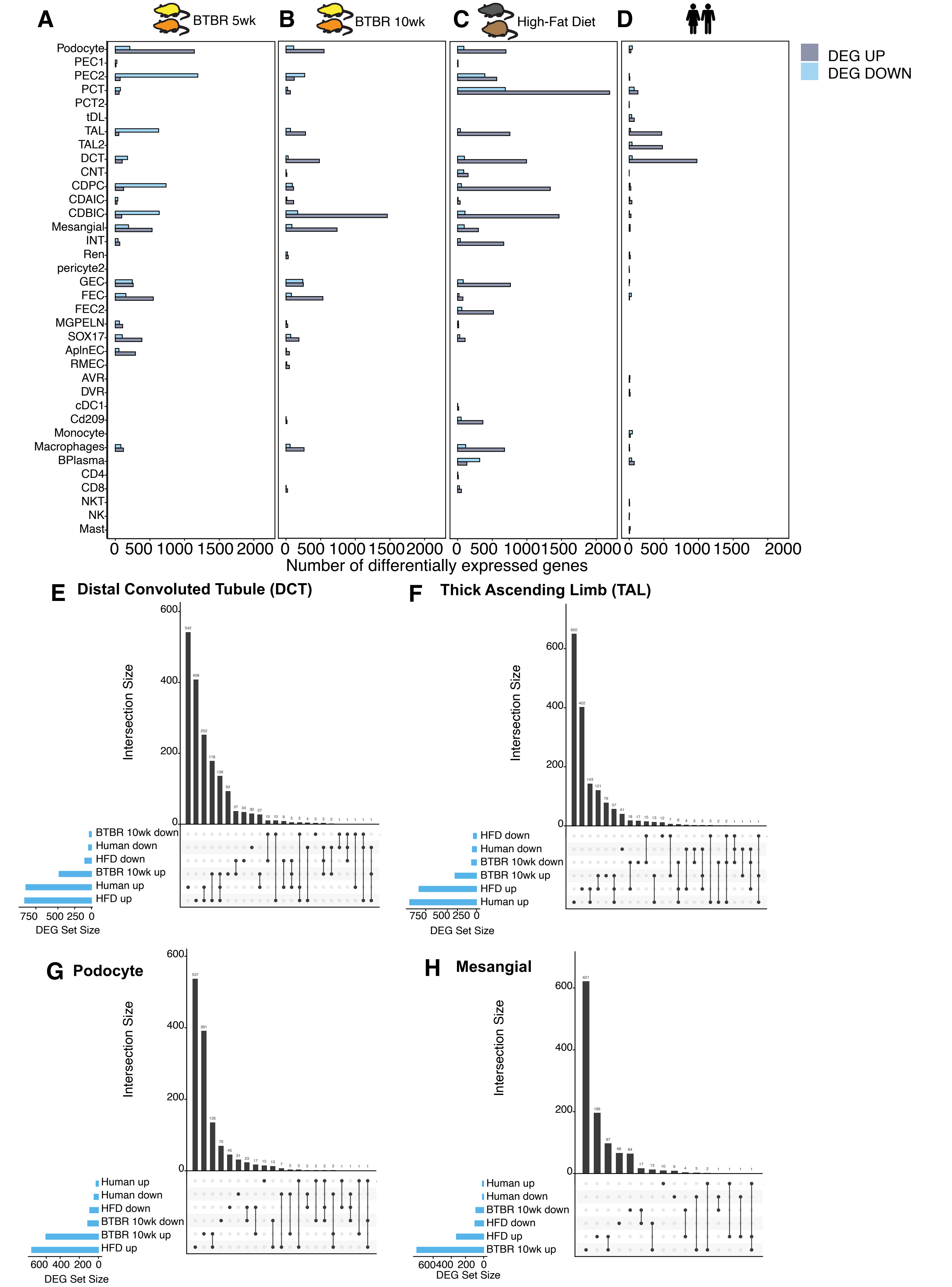

Supplementary Figure 10

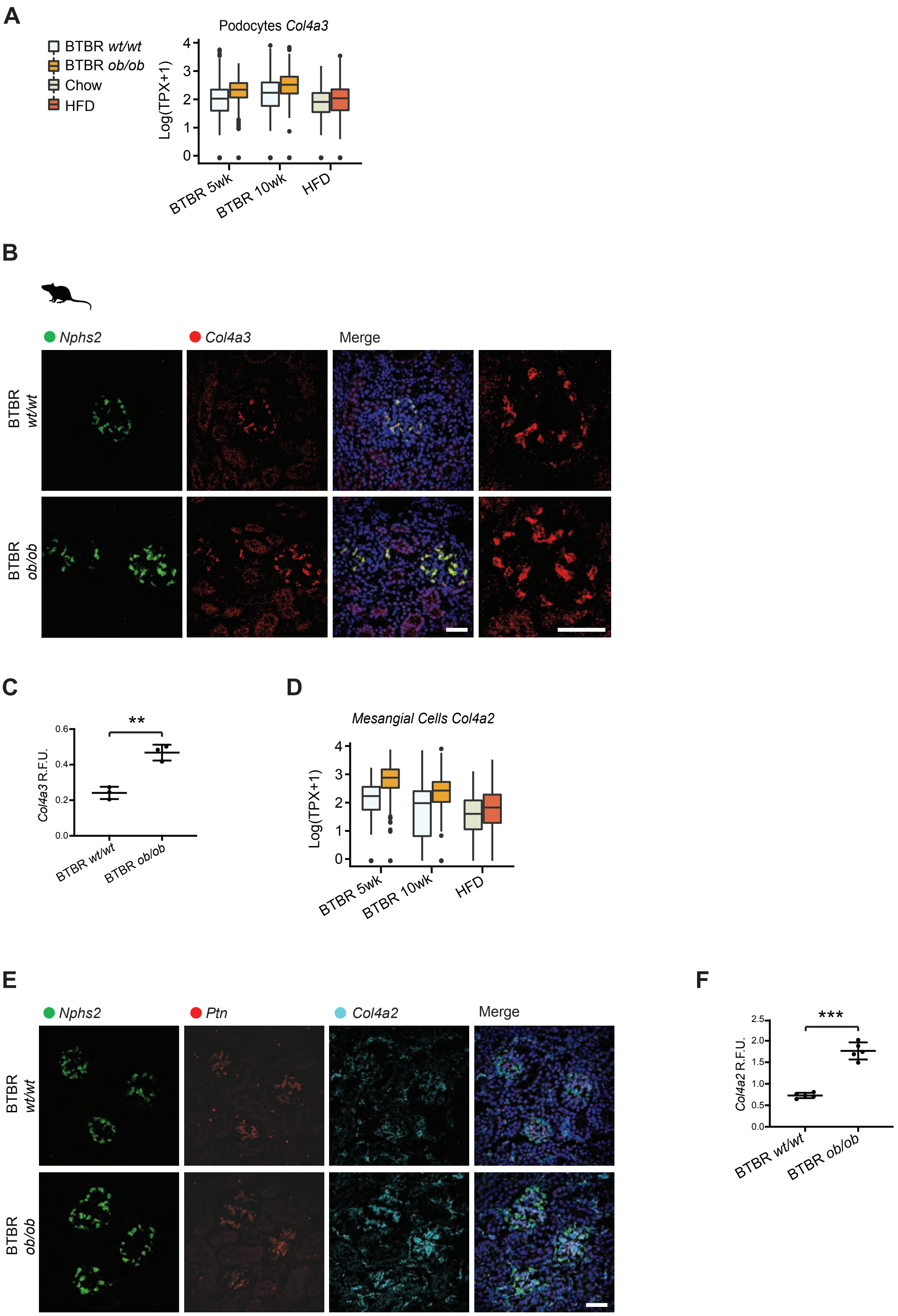

Supplementary Figure 11

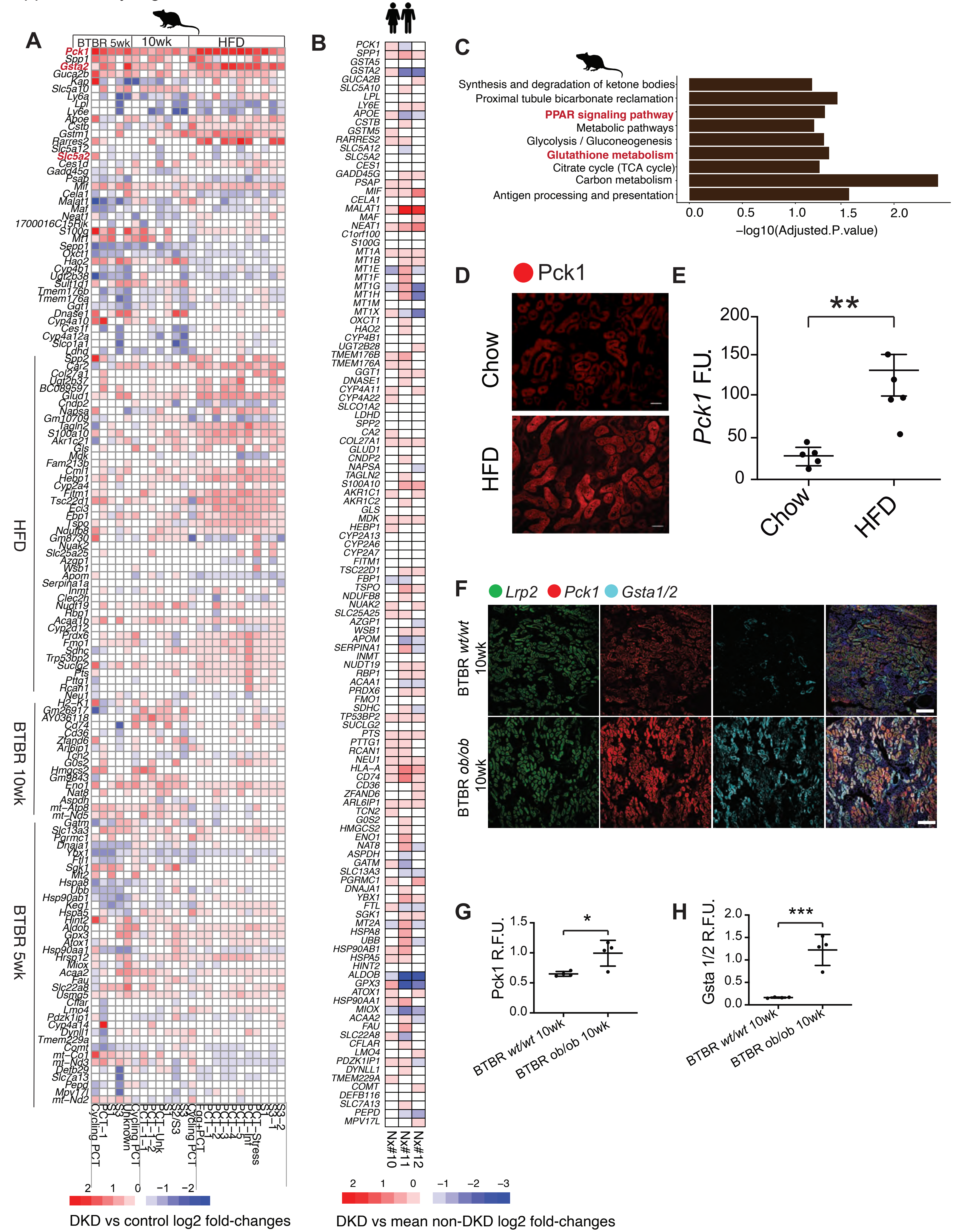

Supplementary Figure 12

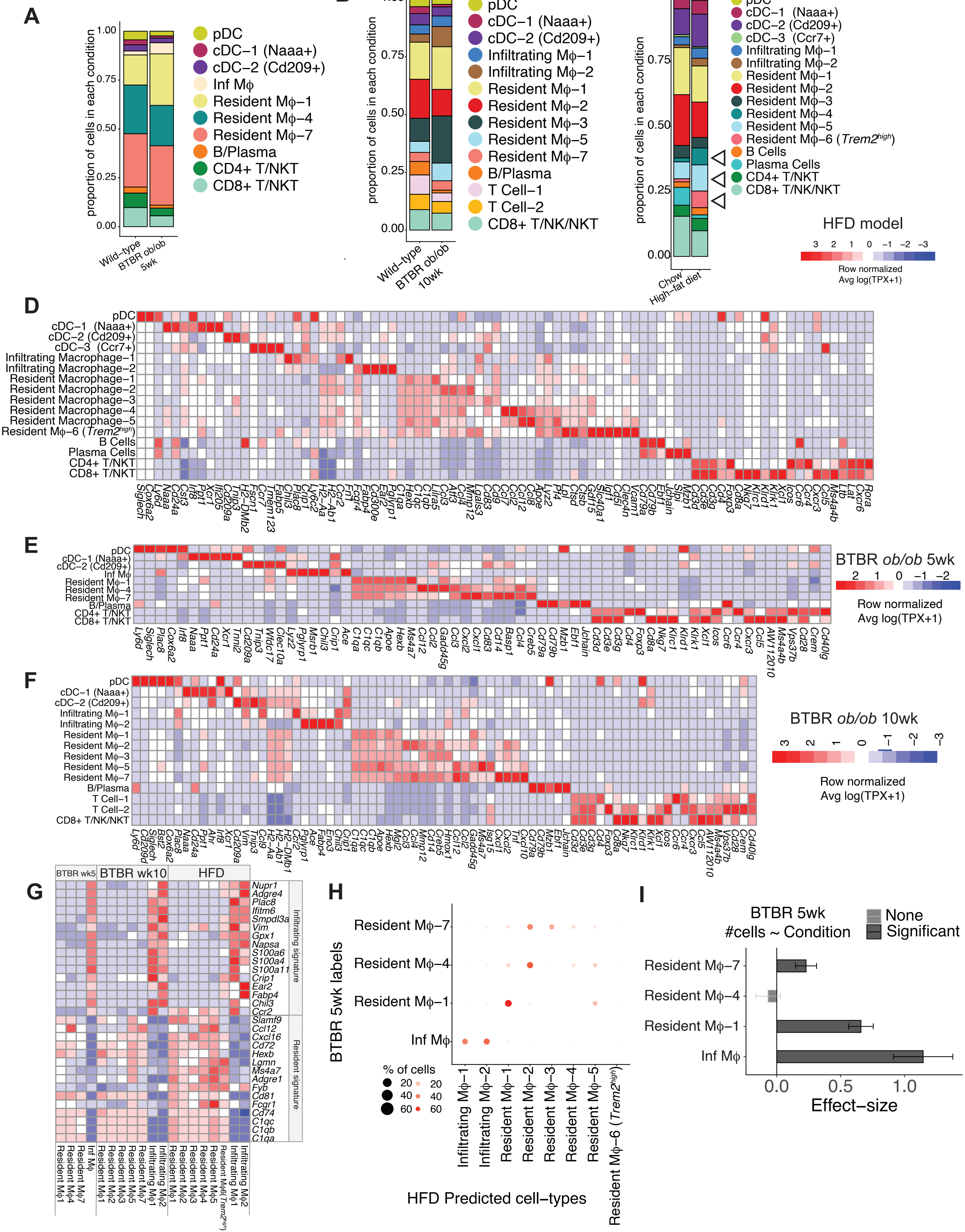
